# Structure of an Archaeal Ribosome with a Divergent Active Site

**DOI:** 10.1101/2024.11.12.623245

**Authors:** Amos J. Nissley, Yekaterina Shulgina, Roan W. Kivimae, Blake E. Downing, Petar I. Penev, Jillian F. Banfield, Dipti D. Nayak, Jamie H. D. Cate

**Affiliations:** Department of Chemistry, University of California, Berkeley, California, United States; Innovative Genomics Institute, University of California, Berkeley, California, United States; Department of Molecular and Cell Biology, University of California, Berkeley, California, United States; California Institute for Quantitative Biosciences, University of California, Berkeley, California, United States; Department of Plant and Microbial Biology, University of California, Berkeley, California, United States; Department of Earth and Planetary Science, University of California, Berkeley, California, United States; Department of Environmental Science, Policy, and Management, University of California, Berkeley, California, United States; Molecular Biophysics and Integrated Bioimaging Division, Lawrence Berkeley National Laboratory, Berkeley, California, United States

## Abstract

The ribosome is the universal translator of the genetic code and is shared across all life. Despite divergence in ribosome structure over the course of evolution, the peptidyl transferase center (PTC), the catalytic site of the ribosome, has been thought to be nearly universally conserved. Here, we identify clades of archaea that have highly divergent ribosomal RNA sequences in the PTC. To understand how these PTC sequences fold, we determined cryo-EM structures of the *Pyrobaculum calidifontis* ribosome. We find that sequence variation leads to the rearrangement of key PTC base triples and differences between archaeal and bacterial ribosomal proteins also enable sequence variation in archaeal PTCs. Finally, we identify a novel archaeal ribosome hibernation factor that differs from known bacterial and eukaryotic hibernation factors and is found in multiple archaeal phyla. Overall, this work identifies factors that regulate ribosome function in archaea and reveals a larger diversity of the most ancient sequences in the ribosome.

## Main

The ribosome is a ubiquitous ribonucleoprotein complex responsible for protein synthesis in the cell. While many aspects of ribosome structure have diverged over the course of evolution, the active sites of the ribosome are thought to be nearly universally conserved^1–3^. The structures of ribosomes from model organisms have been well studied by X-ray crystallography^4–8^, but less is known about how ribosomes differ in non-model organisms. Cryogenic electron microscopy (cryo-EM) has recently enabled the determination of ribosome structures from a wide array of organisms^9–13^ and organelles^14–16^ at high resolution. These structures demonstrate that ribosomes vary in structure and provide a more holistic understanding of ribosome function and regulation.

Archaea are prokaryotes that are widespread on Earth, found in diverse environments, and play important roles in biogeochemical cycling^17–20^. For example, methanogenic archaea are the primary driver of global methane emissions^21^. Some archaea are also hyperthermophiles and have adapted to live in environments that reach extreme temperatures. Hyperthermophiles have modified their translation machinery, including the ribosome, to function at high temperatures. These adaptions include post-transcriptional RNA modifications^22,23^ and changes in ribosomal proteins (rProteins) and ribosomal RNA (rRNA) sequences^24,25^. However, as there are only high-resolution ribosome structures from a few archaeal species^5,22,26–28^, we lack a clear understanding of how archaeal ribosomes differ in their structure and regulation.

In the large subunit (LSU) of the ribosome, the peptidyl transferase center (PTC) catalyzes peptide bond formation during translation. Due to its functional importance, the PTC, which is comprised of rRNA, is one of the most ancient and conserved regions of the ribosome^2,29,30^. Extreme sequence conservation hampers our understanding of how natural changes to the PTC affect its structure and function. While the sequence of the PTC is broadly conserved in archaea, it was previously shown that the ribosomes from *Thermoproteus tenax*, a member of the archaeal class Thermoprotei, have rare sequence variation at rRNA positions in the PTC that are otherwise highly conserved^31^. Here, we find that PTC sequence variation more broadly occurs in a subset of organisms from the archaeal classes Thermoprotei and Methanomethylicia. *Pyrobaculum calidifontis*, an aerobic hyperthermophile (T_opt_=90-95 °C)^32^, is a member of Thermoprotei and has rare variation at five rRNA positions in the core of the PTC. Using cryo-EM, we determine the structure of the *P. calidifontis* ribosome, which reveals how nucleotide changes are accommodated in an archaeal PTC. Furthermore, we show that differences between bacterial and archaeal rProtein uL3 enable greater sequence variation in archaeal ribosomes.

In addition to identifying divergent ribosomal features, we find a protein of previously unknown function bound to the *P. calidifontis* ribosome. We designate this protein **D**ual **R**ibosomal subunit **I**nhibitor (Dri). Dri is comprised of two ribosome binding modules that interact with the LSU and small ribosomal subunit (SSU), where it occupies the PTC and mRNA binding channel. We show that Dri inhibits protein synthesis, consistent with its role as a ribosome hibernation factor. Ribosome hibernation factors play a critical role in preserving ribosomes when translation is downregulated during cellular dormancy, an important part of the microbial life cycle^33,34^. In response to stress or dormancy, ribosome hibernation factors bind to the ribosome, inhibit protein synthesis, and can protect ribosomes from damage while in an inactive state^35–37^. Dri is structurally distinct from known bacterial and eukaryotic hibernation factors and is unique in its ability to interact with both ribosomal subunits independently. We further find that a Dri homolog co-purifies with the ribosomes from a stationary phase culture of *Methanosarcina acetivorans*, a member of the archaeal phylum Halobacteriota. Dri homologs are found broadly in multiple archaeal phyla and may constitute a novel and widely distributed family of archaeal ribosome hibernation factors.

## Results

### Cryo-EM structures of the P. calidifontis ribosome

To investigate the unique features of Thermoproteota (formerly Crenarchaeota) ribosomes, we grew and isolated ribosomes from *P. calidifontis*. 70S ribosomes isolated from archaea in Thermoproteota often dissociate into subunits during ultracentrifugation, even under high magnesium concentrations^28,38,39^. However, while most *P. calidifontis* ribosomes dissociate during ultracentrifugation, some 70S ribosomes stay associated (Extended Data Fig. 1a), as observed for select organisms in Thermoproteota^40,41^. We determined the cryo-EM structure of the *P. calidifontis* 70S ribosome to a global resolution of 2.4 Å (Fig.1a, Extended Fig 2b., and Supplemental Fig. 1). Due to the large amount of dissociated 50S subunits in the sucrose gradient, we also found a population of 50S subunits during cryo-EM processing and determined the *P. calidifontis* 50S subunit structure to a resolution of 2.0 Å.

In the cryo-EM structure of the *P. calidifontis* 70S ribosome, we were able to model all three rRNAs and 61 annotated rProteins (Fig. 1b). Additionally, we found density for four proteins not annotated as rProteins in the *P. calidifontis* proteome. One of the proteins was previously identified as an archaea-specific rProtein, aS21^42^, however homologs were not identified in *Pyrobaculum*. For the other proteins, the cryo-EM maps had sufficient resolution to identify them using the automated modeling software ModelAngelo^43^. Demonstrating the strength of this technique, we were able to identify a protein that was absent from the *P. calidifontis* UniProt proteome (aS35 in Fig. 1b). The other two novel rProteins were annotated as proteins with unknown function, and we designate them as aL48 and aL49. We further confirmed the identities of these proteins using mass spectrometry (Supplemental Tables 1 and 2). These proteins are archaea-specific rProteins that are distributed in the order Thermoproteales (Extended Data Fig 1b.)

**Fig. 1.**
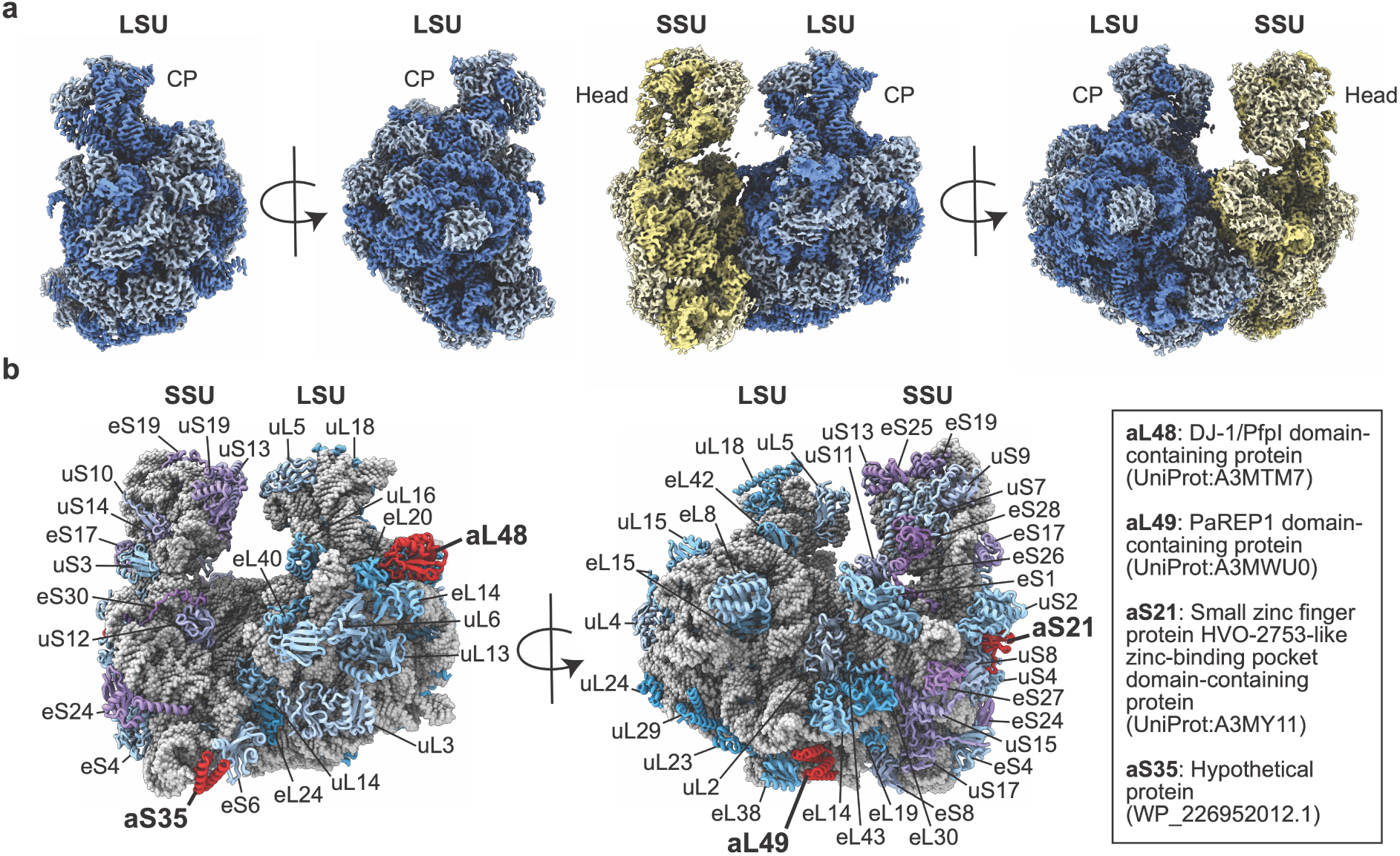
Cryo-EM structures of the *P. calidifontis* ribosome. **a**, Cryo-EM maps for the 50S subunit (LSU, left) and 70S ribosome (right) from *P. calidifontis*. The LSU central protuberance (CP) and SSU head are marked for reference. **b**, rProteins are labeled on the model of the *P. calidifontis* 70S ribosome. Archaea-specific rProteins are shown in red, and their previous annotations, including accession codes, are shown on the right.

Our high-resolution cryo-EM maps also allowed us to identify magnesium and zinc binding sites. The number of zinc binding sites in rProteins was previously shown to correlate with growth temperature^24^. Consistent with this proposal, we identified 11 zinc binding sites in the *P. calidifontis* ribosome (Extended Data Fig. 1c). Next, we identified putative pseudouridylation, nucleobase methylation and acetylation, and 2′-*O*-methylation sites in the *P. calidifontis* ribosome (Extended Data Fig. 1d-e). Previously, it was shown that *N*^4^-acetylcytidine is a widespread modification in ribosomes from the hyperthermophilic archaea *Thermococcus kodakarensis* and that the number of *N*^4^-acetylcytidine modifications increases with growth temperature^22^. In contrast, we find very few *N*^4^-acetylcytidine sites in the *P. calidifontis* ribosome, where the predominant post-transcriptional modification is 2′-*O*-methylation. Additionally, post-transcriptional modifications are mainly buried in the core of the *P. calidifontis* ribosomal subunits and are less widely distributed compared to their distribution in ribosomes from *T. kodakarensis* (Extended Data Fig. 1f). The limited distribution of post-transcriptional modifications has also been observed in the SSU from *Saccharolobus solfataricus*^44^, another member of Thermoprotei.

### Highly divergent peptidyl transferase centers in hyperthermophilic archaea

We aligned 23S rRNA sequences of archaeal species from the Genome Taxonomy Database (GTDB)^45^ and found that certain organisms in the classes Thermoprotei and Methanomethylicia have sequence variation at up to six positions in the PTC. The sequence variation spans the regions 2447-2453 and 2500-2504 (*Escherichia coli* 23S rRNA numbering) at positions that are highly conserved in bacteria and archaea (Fig. 2a). The *P. calidifontis* ribosome has sequence variation at 5 of these 6 conserved positions in the PTC (Fig. 2b). Additionally, we find that many core PTC nucleotides are more conserved across bacteria than archaea, as calculated by Shannon entropy of nucleotide frequencies from 23S rRNA alignments of GTDB species (Extended Data Fig 2a).

**Fig. 2.**
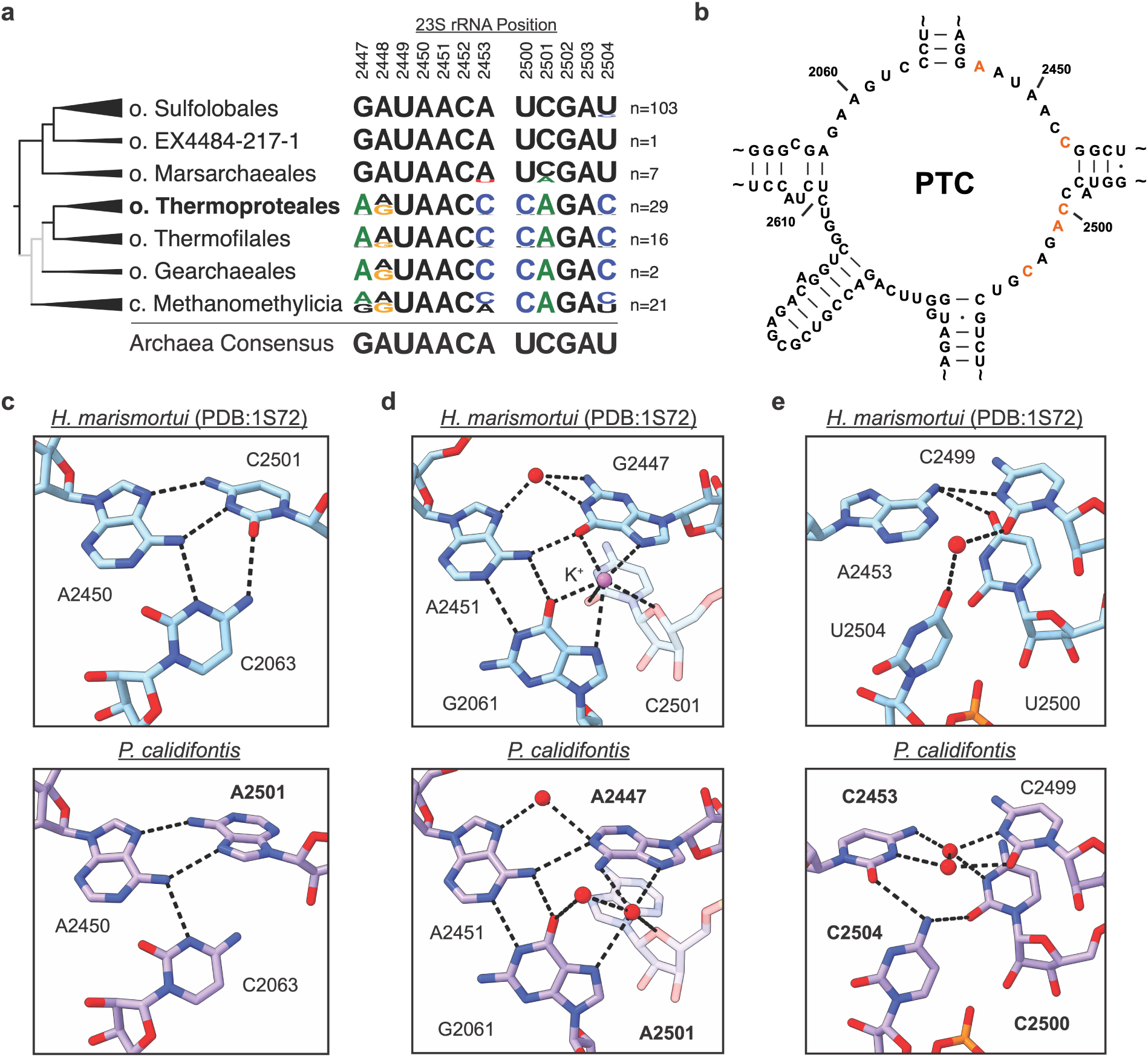
Sequence diversity in archaeal PTCs. **a**, Distribution of rare PTC sequences in archaea. The cladogram is derived from the GTDB phylogeny, where branches with bootstrap support <50% are colored in grey. o and c represent order and class respectively, and n represents the number of 23S rRNA sequences in each distribution. Nucleotides that differ from the archaeal consensus sequence are shown in color. **b**, Secondary structure of the *P. calidifontis* PTC. Positions that differ from the consensus sequence are colored orange. Positions are labeled according to *E. coli* 23S rRNA numbering. **c-e**, Comparison of the PTCs in *H. marismortui* PDB:1S72^46^ (top) and *P. calidifontis* (bottom). Water molecules and potassium ions are shown as red and pink spheres respectively. Hydrogen bonds shown with dashes were calculated in ChimeraX^81^.

We compared the cryo-EM structure of the *P. calidifontis* 50S subunit to the X-ray crystal structure of the *Haloarcula marismortui* 50S subunit (PDB:1S72)^46^, which has a canonical PTC sequence, to understand how nucleotide changes affect the structure of the PTC. Here, we use *E. coli* 23S rRNA numbering and a conversion table to *P. calidifontis* numbering is in Supplemental Table 3. One set of nucleotides affected by rRNA sequence variation is 2063, 2450, and 2501, which forms the canonical C2063-A2450-C2501 base triple in *H. marismortui*. However, in *P. calidifontis* the base triple is disrupted by a C to A change at position 2501. To compensate for this change, A2501 adopts a *syn*-conformation to make a Hoogsteen base pair with A2450 (Fig. 2c and Extended Data Fig. 2b). A second region of the PTC affected by sequence variation in *P. calidifontis* involves the interaction between bases 2061, 2447, and 2451. In *H. marismortui*, G2447 hydrogen bonds with nucleotide A2451 and coordinates a potassium ion that interacts with G2061 and C2501. *P. calidifontis* contains a G2447A change that preserves hydrogen bonding to A2451 (Fig. 2d and Extended Data Fig. 2c), consistent with a report that a G2447A mutation is viable in *E. coli* ribosomes, and ribosomes with this mutation support cellular protein synthesis^47^. However, the A2447 change, along with the *syn* conformation of A2501, results in loss of the potassium ion binding site seen in the *H. marismortui* ribosome, with a concomitant positioning of two water molecules that bridge bases A2061, A2447, and A2501 (Fig. 2d). The third major rearrangement in the PTC occurs due to three differences in the *P. calidifontis* rRNA at positions 2453, 2500, and 2504, which are all cytidines. In *H. marismortui*, these three bases interact with the base of C2499, forming sparse hydrogen bonding. In *P. calidifontis*, these four bases form a more complex hydrogen bonding network that interacts with two ordered water molecules (Fig. 2e and Extended Data Fig. 2d).

A fourth difference in the rRNA from *P. calidifontis* occurs in the A loop of the LSU, which serves as the binding site for the CCA-end of the A-site transfer RNA (tRNA)^48^. We previously reported that rare uridine to cytidine sequence variation at positions 2554 and 2555 of the A loop is found in Thermoprotei archaea and this sequence variation stabilizes the LSU from thermal denaturation^25^. Comparing the A loop from *P. calidifontis* and *H. marismortui*, which has uridines at positions 2554 and 2555, cytidines 2554 and 2555 change positions so that they stack with each other and G2553 (Extended Data Fig. 2e-g). We did not observe this conformational change in *E. coli* ribosomes with U2554C/U2555C mutations that contained an A-site tRNA, suggesting that this conformation may only exist in the apo ribosome^25^. These stacking interactions likely rigidify the A loop, which is part of the last region of the 23S rRNA to fold during ribosome assembly^49^, to help stabilize the LSU.

### Differences in rProtein uL3 enable PTC sequence variation in archaea

*P. calidifontis* PTC sequence variation occurs at rRNA positions where mutations are not tolerated in the *E. coli* ribosome. Specifically, single point mutations at positions A2453C, U2500C, and U2504C inactivate or greatly decrease the activity of *E. coli* ribosomes^50^ and *E. coli* ribosomes with these individual mutations are unable to support growth *in vivo*^51^. Conversely, ribosomes from halophilic archaea with these point mutations can support growth *in vivo*, albeit with decreased growth rates^52,53^. It is unclear why these mutations can be accommodated in archaeal ribosomes while they are not tolerated in *E. coli* ribosomes.

As the *P. calidifontis* ribosome has cytidine residues at positions 2453, 2500, and 2504, we compared its structure to the *E. coli* ribosome^54^ to understand structural differences that enable sequence variation at these positions. In the *E. coli* ribosome, Ψ2504 base pairs with C2452, and A2453 base pairs with U2500, distinct from the complex hydrogen bonding network seen in *P. calidifontis* (Fig. 2e, Fig. 3a). For the interactions to form in *P. calidifontis*, C2504 flips up 54° compared to Ψ2504 in *E. coli*, and to avoid steric clashes, A2572 moves out of the way compared to its position in *E. coli*. However, the shift in A2572 in *P. calidifontis* can only occur because rProtein uL3 is shorter in *P. calidifontis* than in *E. coli* near the PTC (Fig. 3b). The region of uL3 proximal to the PTC is extended in some bacteria compared to archaea and eukaryotes (Extended Data Fig. 3a). If uL3 were not shorter in *P. calidifontis*, A2572 would clash with *E. coli* uL3 residue Q150, which is post-translationally methylated^55^. Interestingly, the flipped conformation of nucleobase 2504 is found in other archaeal ribosomes and is not exclusive to ribosomes with sequence variation at position 2504 (Extended Data Fig. 3c).

**Fig. 3.**
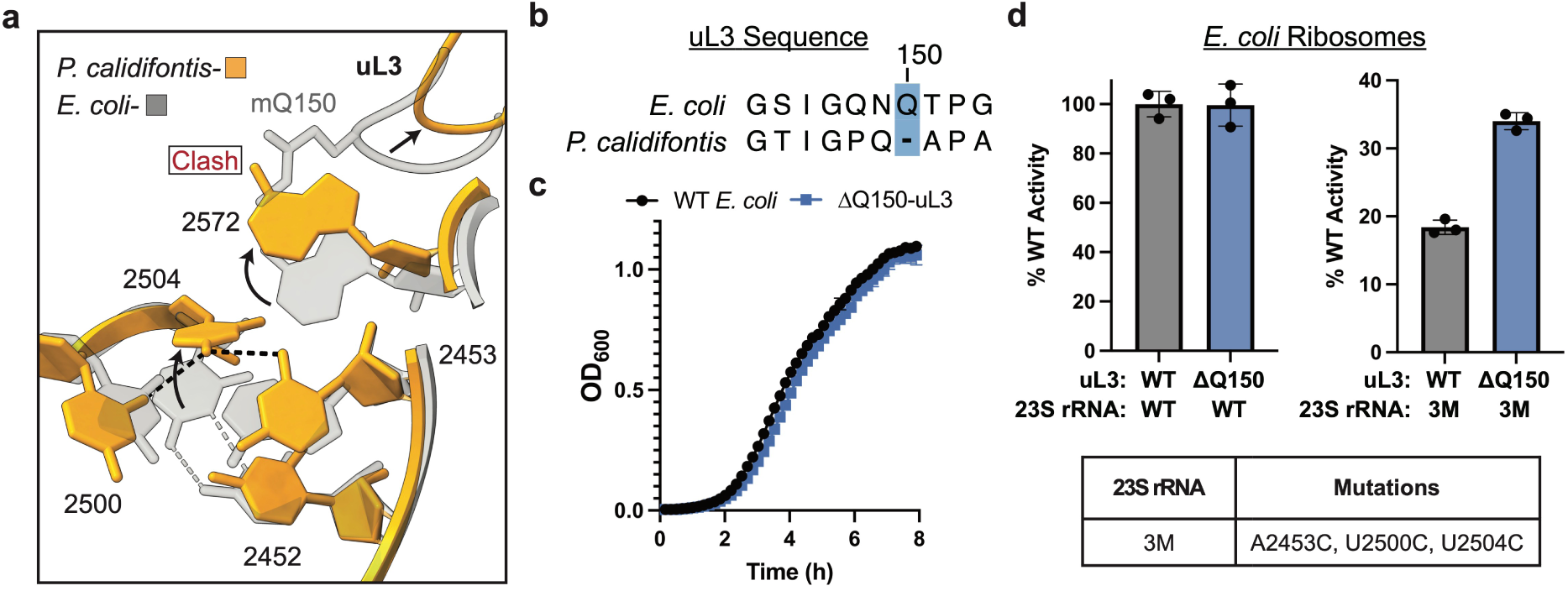
uL3 length affects the mutability of PTC rRNA. **a**, Comparison of the PTC and uL3 in *P. calidifontis* (yellow) and *E. coli* PDB:7K00^54^ (grey). Positions are labeled according to *E. coli* 23S rRNA numbering. **b**, Protein alignment for *E. coli* and *P. calidifontis* uL3 loops that interact with the PTC. **c**, Growth curve of WT and ΔQ150-uL3 *E. coli*. Each data point is the average of three independent measurements and error bars represent the standard deviation of the three measurements. **d**, HiBiT *in vitro* activity assays for mutant *E. coli* ribosomes. Ribosome activity is the slope of the increase in luminescence signal in each *in vitro* translation reaction (methods) and activities were normalized to WT. Each point on the graph represents an individual measurement, and error bars represent the standard deviation of the three measurements.

We wondered whether positions 2453, 2500, and 2504 are mutable in archaea, but not in *E. coli*, because of the rearrangement of base 2504 and a shorter uL3 sequence. To test this hypothesis, we used CRISPR-Cas9 editing^56^ to delete Q150 from the *rplC* gene (ΔQ150-uL3) in *E. coli* (Extended Data Fig. 3b). We find that ΔQ150-uL3 *E. coli* are viable and have no major growth defects other than a modest increase in the lag phase (Fig. 3c). Next, we isolated ribosomes from WT or ΔQ150-uL3 *E. coli* strains using an MS2-tag in helix H98 of the 23S rRNA^57^. *E. coli* ribosomes with ΔQ150-uL3 display WT levels of activity in an *in vitro* translation assay, indicating that the extended uL3 loop is not essential for PTC function. Surprisingly, *E. coli* ribosomes with WT-uL3 and A2453C, U2500C, and U2504C (3M) 23S rRNA mutations have 18% of WT activity, suggesting there is a synergistic effect between these mutations even with the longer uL3 loop. However, consistent with our hypothesis, *E. coli* ribosomes with ΔQ150-uL3 and 3M mutations have 34% of WT activity, demonstrating that the length of uL3 modulates the mutability of these rRNA positions.

### Dri is a novel archaeal ribosome hibernation factor

In the *P. calidifontis* cryo-EM maps, we found density in both the isolated 50S subunit and the 70S complex that did not correspond to a known component of the ribosome (Fig. 4a). After 3D classification (Supplemental Fig. 2), we were able to attribute the density to a protein of unknown function using the protein sequence predicted by ModelAngelo^43^ and we verified its presence by mass spectrometry (Supplemental Tables 1 and 2). We designate this protein **D**ual **R**ibosomal subunit **I**nhibitor (Dri) due to its positioning in the ribosome. Dri contains eight cystathionine-β-synthase (CBS) domains^58^ that form two disc-shaped lobes, each comprised of four CBS domains (Fig. 4a). We found that the Dri C-terminal lobe binds to the P and E sites of the SSU on the 70S ribosome (Fig. 4b) and interacts with the ribosome in the rotated state (Extended Data Fig. 4a). An extended loop of the C-terminal lobe reaches into the SSU and blocks the mRNA binding channel (Fig. 4b). Additionally, there is a disordered 24 amino acid loop in the C-terminal lobe that points towards the A site and could block the binding of the A-site tRNA in the SSU. In the majority of 70S particles, the Dri C-terminal lobe is bound to the SSU. However, we also see populations of 70S particles where the N-terminal lobe is bound to the LSU and where both the N and C-terminal lobes are bound to the ribosome concurrently (Supplemental Fig. 2).

**Fig. 4.**
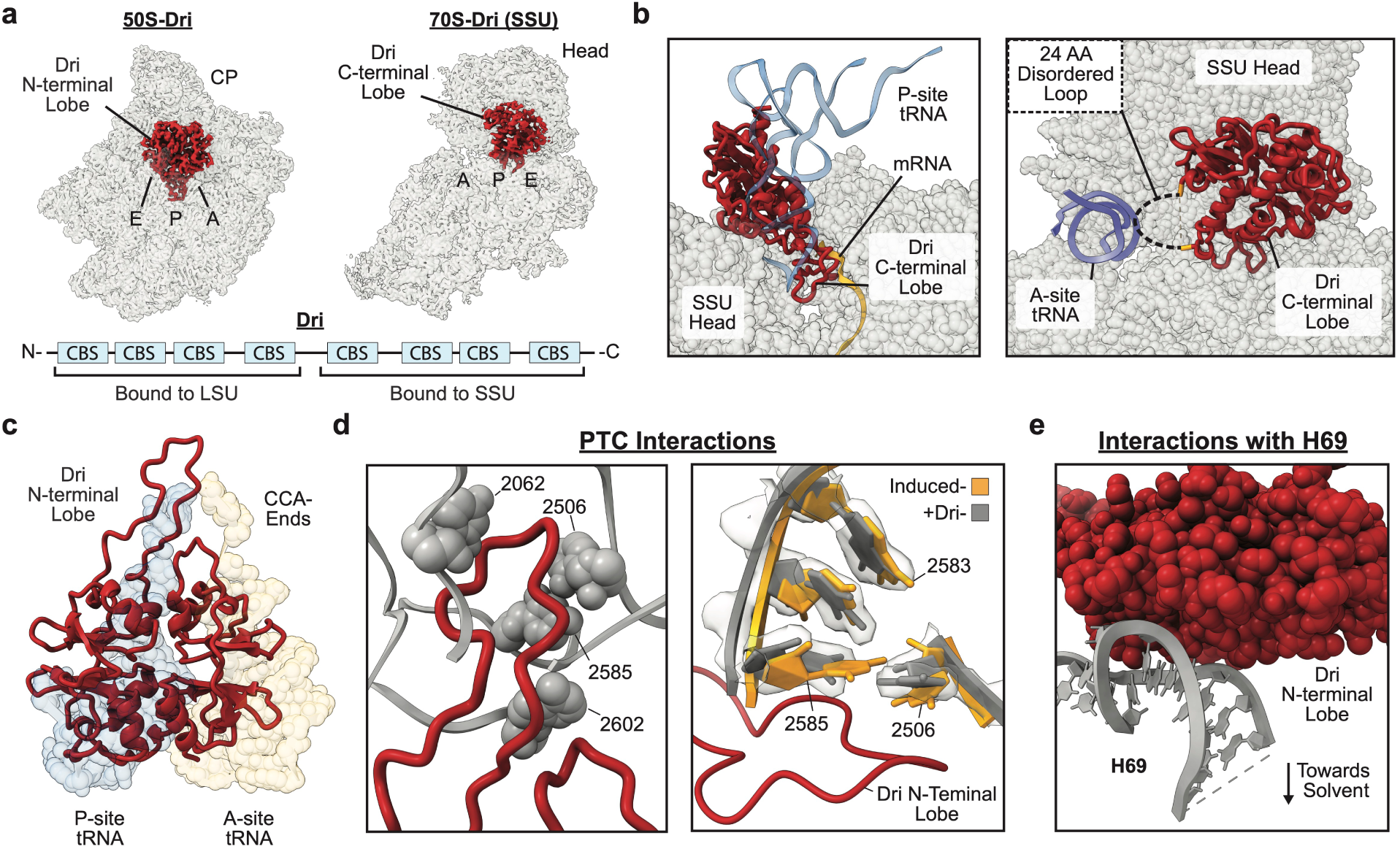
Dri binding sites on the ribosome. **a**, Cryo-EM density for the N-and C-terminal lobes of Dri on the isolated 50S (left) and SSU of the 70S ribosome (right) viewed from the subunit interface. The SSU head and LSU central protuberance (CP) are labeled. The domain architecture of Dri is shown below. **b**, (left) The Dri C-terminal lobe binds to the small subunit and blocks the mRNA channel in the P and E sites. The mRNA and P-site tRNA from PDB:7K00^54^ are overlayed on the *P. calidifontis* structure. (right) A disordered loop of Dri points into the A site of the SSU. The A-site tRNA from PDB:7K00^54^ is overlayed on the *P. calidifontis* structure. **c**, The Dri N-terminal lobe occupies a similar position to the A– and P-site tRNAs on the LSU from PDB:7K00^54^. **d**, (left) The P-site loop of Dri extends into the PTC and contacts key PTC nucleotides. (right) PTC nucleotides that participate in the tRNA induced fit are shown when tRNA is bound to the *Thermus thermophilus* ribosome (gold, PDB:1VY4^62^) or when Dri is bound to the *P. calidifontis* LSU (grey and cryo-EM density). **e**, The N-terminal lobe of Dri makes extensive contacts with H69 of the LSU.

The Dri N-terminal lobe is bound to the isolated 50S subunit and the LSU of the 70S ribosome (Supplemental Fig. 2) and occupies the A-site and P-site tRNA binding sites (Fig. 4c). There is an extended protein loop off the N-terminal lobe that reaches into the P-site of the PTC and interacts with core PTC nucleotides. Interestingly, the N-terminal lobe mimics tRNA binding to the LSU in multiple ways. First, Dri binding stabilizes uL16 in an ordered conformation previously observed upon tRNA binding in the P site ^59,60^ (Extended Data Fig. 4c). Dri also occupies the positions of the A– and P-site amino acids substrates^61^ and further interacts with the A-site cleft^62^, the binding site for the A-site amino acid (Extended Data Fig. 4c). Finally, binding of Dri to the LSU induces an active conformation of the PTC seen when tRNA substrates are bound in both the A and P sites^63^. (Fig. 4d). There is strong density for PTC nucleotides when the Dri N-terminal lobe is bound to the 50S subunit (Extended Data Fig. 4d), while in its absence, several regions of the PTC are disordered. Dri further interacts with the 50S subunit by making extensive contacts with helix H69, a region of 23S rRNA that makes important bridges with the SSU during subunit joining^64^.

Since Dri occludes the tRNA binding sites on the LSU and SSU we hypothesized that Dri is an archaeal-specific ribosome hibernation factor. We developed a *P. calidifontis* lysate-based *in vitro* translation system to translate a HiBiT peptide reporter^65^ using the optimized conditions for hyperthermophilic archaeal *in vitro* translation systems^40^. Upon addition of an mRNA encoding the HiBiT peptide to the *in vitro* reaction, we observed robust translation at 95 °C (Fig. 5a). We recombinantly expressed the full-length WT Dri protein and the isolated Dri N-terminal and C-terminal lobes (Extended data Fig. 5a). Addition of full-length Dri to the *in vitro* translation reaction inhibited peptide output substantially, as did addition of the isolated N-terminal and C-terminal lobes (Fig. 5a). Taken together, these results show that each lobe of Dri can inhibit translation in isolation.

**Fig. 5.**
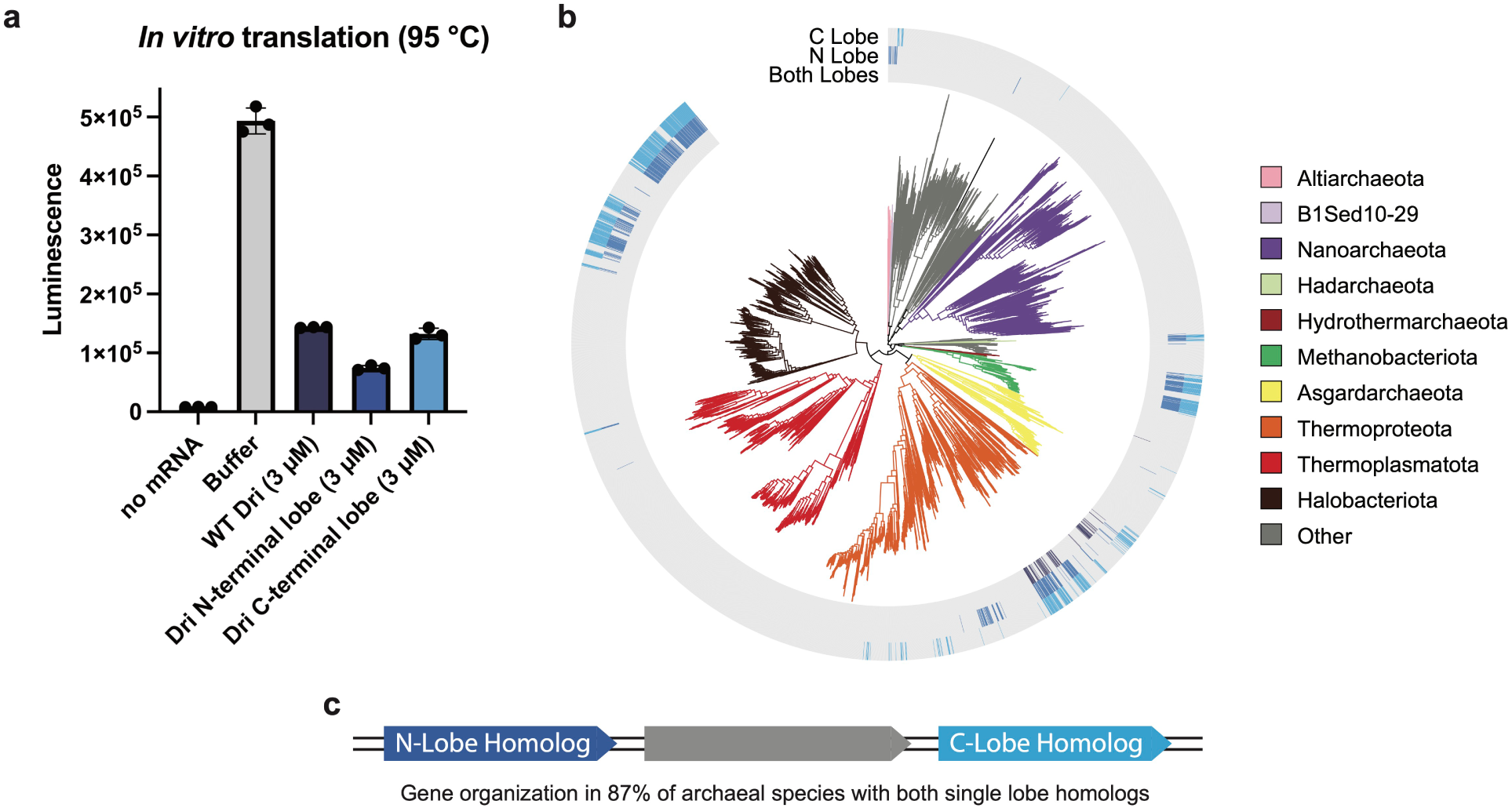
Translation inhibition and phylogenetic distribution of Dri homologs. **a**, *P. calidifontis* lysate-based *in vitro* translation assay. After translation of the HiBiT peptide, luminescence was measured from the HiBiT-LgBiT complex. For the no mRNA and buffer controls, 3 µL of Dri buffer A was added to the reaction. Each point on the graph represents an individual measurement, and error bars represent the standard deviation of the three measurements. **b**, Phylogenetic distribution of Dri homologs indicated on the GTDB archaeal phylogeny. Dri homologs with only a C-terminal or N-terminal lobe are shaded in the outer and middle rings. Dri homologs with two lobes that align to both Dri lobes are shaded in the inner ring. **c**, Archaeal genome architecture of Dri homologs.

We wondered if Dri is broadly present in Archaea or represents a more specialized ribosome hibernation factor for hyperthermophilic archaea. We searched the genomes of all GTDB archaeal species for Dri homologs by sequence homology, which were confirmed to have sequence features used by Dri to bind to the ribosome. We found full length homologs (containing both the N– and C-terminal lobes) mainly in the archaeal phylum Thermoproteota (Fig. 5b and Extended Data Table 1). These homologs contain extended loops in both the N– and C-terminal lobes that likely interact with the PTC and mRNA channel in a similar manner to Dri. Interestingly, we also found truncated Dri homologs comprised of only the N– or C-terminal lobe broadly distributed in at least seven archaeal phyla including Halobacteriota and Methanobacteriota. As the isolated Dri N– and C-terminal lobes inhibit *P. calidifontis* translation, we hypothesize that single-lobed Dri homologs function as ribosome hibernation factors. The single-lobe homologs frequently co-occur, as 59% of genomes that contain one homolog also contain the other, and they are typically found in the same gene order with an additional gene interspersed (Fig. 5c). To further asses if single-lobed Dri homologs associate with the ribosome, we isolated 70S ribosomes from a stationary phase culture of *M. acetivorans*, a member of Halobacteriota (Extended Data Fig. 5b). We detected the *M. acetivorans* C-terminal lobe homolog in the 70S ribosome sample with mass spectrometry, suggesting that this homolog also binds to the ribosome (Supplemental Table 4). Interestingly, we also found the *M. acetivorans* CBS-domain containing protein that corresponds to the gene interspersed between the N– and C-terminal lobe homologs in the 70S ribosome sample, consistent with these proteins binding to ribosomes.

## Discussion

Here, we show that one of the most conserved elements of the ribosome, the PTC, is highly divergent in certain archaeal ribosomes. Organisms in the archaeal classes Thermoprotei and Methanomethylicia harbor sequence variations in the core of the PTC at up to six positions, a region that is highly conserved in bacteria. Using cryo-EM, we determined the structure of the Thermoprotei *P. calidifontis* ribosome, revealing how sequence variation in the PTC results in alternative structural arrangements that position the non-varying nucleotides in the correct position seen in other ribosomes. While it remains unclear why these branches of Archaea have a high level of PTC sequence variation, four of the nucleotide changes found in Thermoprotei and Methanomethylicia were shown to confer resistance to PTC inhibitors, including linezolid^53^ (A2453C, U2500C, and U2504C), anisomycin^52,66^ (G2447A, A2453C, and U2500C), and chloramphenicol^67^ (U2504C). This suggests that one consequence of PTC variation could be the evasion of PTC inhibition by small molecules.

PTC sequence variation occurs at positions in the *P. calidifontis* ribosome that cannot be mutated in ribosomes from the model bacterium *E. coli*^50,51^. Our finding that natural PTC sequence variation is more widespread in Archaea than in Bacteria could suggest that there are differences in bacterial ribosomes that limit the variability of the PTC compared to archaea. We hypothesize that one of these is the shorter length of rProtein uL3 in archaea compared to bacteria. To mimic archaeal uL3 sequences, we delete a single amino acid in *E. coli* uL3, which has little effect on *E. coli* growth and ribosome function. We find that the shorter uL3 increases the activity of *E. coli* ribosomes with A2453C, U2500C, and U2504C mutations, suggesting that differences in the mutability of PTC positions in *E. coli* and archaea could be due to PTC rearrangement near uL3. In further support of this, it was shown that changes in the uL3 sequence proximal to the PTC affects the conformation of PTC nucleotides in *Saccharomyces cerevisiae*^68^. The difficulty of mutating multiple PTC positions in *E. coli* has been a challenge in efforts to engineer the PTC for novel functions^69–71^. However, it has been shown that epistatic interactions in the PTC can compensate for deleterious mutations^72^. Here, we show that uL3 sequence and length modulate the mutability of PTC positions and demonstrate that modified rProteins could aid in the development of ribosomes with engineered active sites.

Ribosome hibernation has been well studied in bacteria and eukaryotes, but it was previously unknown what factors influence hibernation in archaea.^36,37^ While a factor that dimerizes isolated 30S subunits was identified in the genus *Pyrococcus*, it does not dimerize 70S ribosomes, blocks 70S formation, and it remains unclear if it participates in ribosome hibernation or plays some unknown role.^73^ Here, we describe a novel archaeal hibernation factor, Dri, that binds to the LSU and SSU of the ribosome and inhibits translation. The Dri C-terminal lobe binds to the mRNA channel in the P and E sites, similar to known bacterial and eukaryotic hibernation factors^74–76^. The independent evolution of hibernation factors that target the mRNA channel in each domain of life underscores the importance of inactivating the SSU during hibernation.^36,37^ In addition, the Dri N-terminal lobe binds to the LSU and makes extensive contacts in the PTC. Certain eukaryotic and bacterial hibernation factors (i.e. MDF2^76^ and Balon^13^, respectively) also interact with the PTC. Dri differs from these proteins in its ability to mimic the binding of tRNAs to the LSU, even causing the PTC to adopt an active conformation, which is surprising for a protein that inhibits translation. This could suggest that it in addition to blocking tRNA binding, the Dri N-terminal lobe helps to chaperone the PTC during hibernation. In line with this hypothesis, we find that the *P. calidifontis* PTC is disordered in the absence of Dri. Interactions between the Dri N-terminal lobe and H69 in the LSU, which makes important contacts with the SSU during subunit joining, could also suggest that the N-terminal lobe may play a role in stabilizing hibernating 70S ribosomes. This could explain why we were able to isolate *P. calidifontis* 70S ribosomes from sucrose gradients, while the ribosomes from some organisms in Thermoproteota, completely dissociate during purification^28,38–41^.

Dri is an archaea-specific ribosome hibernation factor that evolved separately from known bacterial and eukaryotic hibernation factors. While some Dri homologs have two ribosome binding lobes, other homologs have a single lobe that would only interact with one of the ribosomal subunits. In many genomes, two single-lobed homologues are found in the same orientation, typically separated by a single gene, suggesting that full length Dri in Thermoproteota may have evolved by gene fusion of the more widely distributed single-lobed gene organization. Since the isolated N and C-terminal Dri lobes can independently inhibit translation, the two single-lobed Dri homologs likely can work together to inhibit translation in a manner similar to the two-lobed homologs. Dri homologs are widespread in archaea and are found broadly in multiple phyla. While it has been suggested that hibernation factors evolved after the split between Bacteria and Archaea/Eukaryota^36^, the discovery of an archaea-specific hibernation factor suggests that ribosome hibernation may have also evolved independently in Archaea and Eukaryotes. Further studies of archaeal ribosome hibernation will help to uncover the evolution of this ubiquitous yet diverse regulatory process.

## Methods

### Pyrobaculum calidifontis growth

*Pyrobaculum calidifontis* (DSM 21063) was purchased from the DSMZ collection and grown as described^32^ with modifications. For long-term storage, cells were kept at –80 °C in 40% (v/v) glycerol. Cells were grown in modified TY medium (0.1% yeast extract, 1% tryptone, 0.3% sodium thiosulphate, pH 7.0) at 90 °C without shaking. For pre-cultures, 200 mL of modified TY media in a 1 L screw top bottle was inoculated with 600 µL of a frozen cell stock. The pre-culture was placed in an oven at 90 °C without shaking for two days. Cultures (200 mL of modified TY medium in a 1 L screw top bottle) were then inoculated with 20 mL of pre-culture and incubated at 90 °C until they reached an OD_600_ of 0.18.

### *P. calidifontis* ribosome preparation

3 L of *P. calidifontis* cell culture at stationary phase (OD_600_ of 0.18) were centrifuged at 6,000 *xg* for 20 minutes at 4 °C. The cell pellet was then resuspended in 20 mL *Pc* buffer A (20 mM HEPES-KOH pH 7.0, 100 mM NH_4_Cl, 10 mM Mg(OAc)_2_, 2 mM dithiothreitol (DTT)). Cells were then lysed by sonication and spun in an F14-14×50cy rotor (ThermoFisher) at 12,000 rpm (25,000 *xg*) for 45 minutes. The supernatant was then split evenly and layered onto two sucrose cushions containing 35 mL *Pc* buffer A with 1 M sucrose. The sucrose cushions were placed in a Ti-45 rotor (Beckman-Coulter) and centrifuged at 38,000 rpm (113,000 *xg*) for 16 hours at 4 °C. The pellets (crude ribosomes) were then resuspended in 1 mL of *Pc* buffer A. After resuspension, 0.5 mg of crude ribosomes were placed on a 15-35% sucrose gradient in *Pc* buffer B (20 mM HEPES-KOH pH 7.0, 10 mM KCl, 10 mM Mg(OAc)_2_, 2 mM DTT). The sucrose gradient was placed in an SW-32 rotor (Beckman-Coulter) and centrifuged at 26,000 rpm (83,000 *xg*) for 16 hours. After centrifugation, the gradient was fractionated with an ISCO fractionation system. The peak corresponding to 70S ribosomes (Extended Data Fig. 1a) was isolated, buffer exchanged into *Pc* buffer C (20 mM HEPES-KOH pH 7.0, 10 mM KCl, 20 mM Mg(OAc)_2_, 3 mM spermine, 2 mM DTT), and concentrated in a 100 kDa molecular weight cut off spin filter (Millipore). Ribosomes were then flash frozen and stored at –80 °C. *P. calidifontis* ribosomes were quantified using the approximation 1 A_260_=40 µg/mL.

### Recombinant expression and purification of Dri

*E. coli* Rosetta (DE3) cells were transformed with a pET-21(+) plasmid containing the Dri protein sequence with an N or C-terminal His-tag (Supplemental Table 5). Overnight cultures of the transformants were diluted 1:100 into 1 L of LB media with 100 μg/mL ampicillin and incubated at 37°C. Once the OD_600_ reached 0.5, protein expression was induced using 0.5 mM Isopropyl β-D-1-thiogalactopyranosie (IPTG) and the culture was incubated at 37 °C for an additional 2 hours. The cells were then pelleted and flash frozen. Cell pellets were thawed on ice and resuspended in Dri lysis buffer (30 mM HEPES-KOH pH 7.5, 500 mM KCl, 20 mM imidazole) with a Pierce protease inhibitor tablet (ThermoFisher) and lysed by sonication. The lysate was heat treated at 90 °C for 30 minutes and clarified by centrifugation at 25,000 *xg* for 1 hour in an F14-14 x 50cy rotor (Thermo Fisher). The supernatant was collected and filtered through a 0.2 μm filter. A 5 mL His-trap column (Cytiva) was equilibrated with 5 column volumes (CV) of Dri lysis buffer, and the lysate was then loaded onto the column. The column was washed with 10 CV of Dri wash buffer (30 mM HEPES-KOH pH 7.5, 20 mM imidazole) with 1.5 M KCl for the Dri C-terminal lobe or Dri wash buffer with 2.5 M KCl for the Dri N-terminal lobe and WT Dri protein. The protein was eluted using a gradient of 0-100% Dri elution buffer (30 mM HEPES-KOH pH 7.5, 500 mM KCl, 500 mM imidazole) over 5 CV. After the elution, the Dri C-terminal lobe was buffer exchanged into Dri buffer A (30 mM HEPES-KOH pH 7.5, 500 mM KCl, 20% glycerol), concentrated in a 10 kDa cut-off spin filter (Millipore), flash frozen, and stored at – 80 °C.

After elution from the His-trap column, the Dri N-terminal lobe and WT Dri protein were buffer exchanged into heparin buffer A (30 mM HEPES-KOH pH 7.5, 500 mM KCl). A 5 mL heparin column (Cytiva) was equilibrated with heparin buffer A, and the protein was loaded onto the column. The column was then washed with five columns of heparin buffer A. The protein was eluted from the column using a gradient from 0-100% of heparin elution buffer (30 mM HEPES-KOH pH 7.5, 2 M KCl) over 10 CV. Fractions containing protein were combined and buffer exchanged into Dri buffer A. The purified proteins were concentrated in 10 kDa cut-off spin filters (Millipore), flash frozen, and stored at –80°C.

### P. calidifontis in vitro translation

*P. calidifontis* was grown in 1.6 L of modified TY medium at 90 °C without shaking until cultures reached an OD_600_ of 0.09. Cultures were then placed in an ice water bath and after cooling, cells were pelleted at 6,000 *xg* for 20 minutes at 4 °C. The cells were resuspended in 10 mL of *Pc* buffer A and transferred to a 50 mL falcon tube. Cells were then pelleted again at 6000 *xg* for 20 minutes, and the cell pellet was weighed. The pellet was resuspended with 2.5 mL *Pc* buffer C without spermine per gram of wet cell pellet. Cells were lysed by sonication for three minutes and the lysate was clarified by two rounds of centrifugation at 20,000 *xg* for 30 minutes. The lysate was then flash frozen and stored in single-use aliquots at –80 °C.

HiBiT peptide mRNA was *in vitro* transcribed with T7 RNA polymerase and purified using the RNA clean and concentrator kit (Zymo). 30 µL *in vitro* translation reactions were assembled as follows: 15 µL *P. calidifontis* lysate, 3 mM ATP, 2 mM GTP, 3 mM spermine, 100 µM amino acid mixture (Promega), 900 ng HiBiT mRNA, and 10 mM MgCl_2_ (final concentration of 20 mM Mg^2+^). Either 3 µL of Dri protein or Dri buffer A (control reactions) was added to *in vitro* translation reactions. Reactions were incubated at 95 °C for 10 minutes and then cooled on ice. For each reaction, 8 µL was placed in three wells of a 384-well plate. 20 µL of LgBiT buffer (20 mM HEPES pH 7.5, 50 mM KCl, and 10% glycerol) with a 1:50 dilution of NanoGlo HiBiT lytic substrate (Promega) and a 1:100 dilution of LgBiT protein (Promega) was added to each well. The plate was incubated for 10 minutes at room temperature and then luminescence was measured in a Spark plate reader (Tecan).

### Cryo-EM sample preparation and data acquisition

A *P. calidifontis* 70S ribosome aliquot was thawed and diluted to 0.12 mg/mL in *Pc* buffer C. The solution was incubated at 80 °C for 20 minutes and placed on ice. Three-hundred mesh R1.2/1.3 UltrAuFoil grids with a layer of 2 nm amorphous carbon (Quantifoil) were glow discharged in a Pelco easiGlow for 12 seconds under a 0.37 mBar vacuum with 25 mA current. 4 µL of ribosome sample was applied to the grid, incubated for one minute at room temperature, and plunge-frozen in liquid ethane using a Vitrobot Mark IV at 4 °C with 100% humidity.

Cryo-EM movies were collected with a 300 kV Titan Krios microscope with a BIO-energy filter and Gatan K3 camera (Supplemental Table 6). Data was collected with a super-resolution pixel size of 0.41 Å (physical pixel size of 0.8293) over a defocus range of –0.5 to –1.5 µm with a total electron dose of 40 e^-^/Å^2^ split over 40 frames. Serial-EM^77^ was used to automate data collection and image shift was used to collect movies in a 10 x 10 grid of holes with two movies collected per hole.

### Cryo-EM image processing

Image processing was performed in cryoSPARC 4^78^ (Supplemental Fig. 2). Movies were Fourier-cropped to the calibrated physical pixel size (0.8293 Å), patch motion corrected, and the CTFs of micrographs were estimated using patch CTF estimation. Micrographs were split into exposure groups based on their position in the 10 x 10 groups during image-shift collection and micrographs with poor estimated CTF fit were manually excluded from further processing.

Particles were picked with the cryoSPARC template picker using ribosome 2D templates generated in cryoSPARC. Particles were extracted and Fourier-cropped to 1/8 of the box size and 2D classification was performed using 75 classes. After selecting classes corresponding to ribosomes, a second round of 2D classification was performed using 50 classes. The selected particles were reextracted and Fourier-cropped to 1/4 of the box size. A reference map was generated with an *ab initio* reconstruction of the binned particles in cryoSPARC. 3D classification was performed using heterogeneous refinement in cryoSPARC with six classes. Classes containing features consistent with 70S and 50S ribosomes were subjected to a second round of 3D classification.

3D classes consistent with 50S ribosomes were sorted into two classes based on Dri occupancy using 3D classification (without alignment) in cryoSPARC. Particles from the 50S class with stronger density for Dri were reextracted at the full box size. Particles were subjected to homogeneous refinement with the per-particle defocus optimization, per-group CTF parameter optimization, and minimization over per-particle scale options selected. Local motion was corrected using reference-based motion correction in cryoSPARC. The corrected particles were subjected to another homogeneous refinement with Ewald sphere correction to yield the final map.

3D classes consistent with 70S ribosomes were sorted into four classes using a mask on the SSU to remove isolated 50S subunits. Classes consistent with 70S ribosomes were pooled and reextracted at the full box size. Particles were subjected to homogeneous refinement with the per-particle defocus optimization, per-group CTF parameter optimization, and minimization over per-particle scale options selected. Local motion was corrected using reference-based motion correction in cryoSPARC, and a consensus 70S ribosome map was generated with homogeneous refinement. Masks were generated for Dri on the LSU, SSU, and both subunits and were used to classify 70S ribosomes based on protein occupancy. Local resolution was calculated in Relion 4^79^ using the Relion implementation. For figure presentation, cryo-EM maps were supersampled using the ‘make a smoother copy’ function in COOT^80^.

### Model Building

For modeling 70S ribosomes, composite maps were generated for the 70S-consensus reconstruction and 70S-SSU-Dri reconstruction. We aligned the focused-refined maps (SSU-focused, LSU-focused, SSU-head-focused, and LSU-CP-focused for the 70S-consensus reconstruction and SSU-focused, LSU-focused, and SSU-head-focused refined maps for the 70S-SSU-Dri reconstruction) to the global 70S ribosome map using the ‘Fit in Map’ tool in ChimeraX^81^. The focus-refined maps were then resampled using ‘vop resample’ and scaled according to their standard deviations from the ‘Map statistics’ tool. The scaled maps were then added sequentially using ‘vop add’.

Initial models were built using ModelAngelo^43^ provided with sequences for *P. calidifontis* rRNAs (NCBI database) and rProteins (Uniprot). Unknown proteins were built using ModelAngelo without provided sequences. Protein models were inspected for completeness and manually built in regions where the sequences were missing or incorrect. RNA models were manually inspected at each nucleotide for correct modeling and post-transcriptional modifications. To model Dri on the SSU, the AlphaFold prediction^82^ was docked into the cryo-EM map and regions of the prediction that did not agree with the map were rebuilt manually.

Magnesium ions and spermine molecules were identified using the ‘unmodeled blobs’ tool in COOT^80^ and modeled manually. No attempt was made to systematically model potassium ions. The models were then refined in Phenix^83^ with real-space refinement, and manual adjustments were made to the model in COOT^80^ (Supplemental Table 7). After refinement, phenix.douse^83^ was used to model solvent using the default settings. For the 70S structures, solvent was modeled into the individual focused refinement maps and then docked into the composite map.

### Bioinformatic analysis

23S rRNA sequences in archaea and bacteria were identified by searching all Genome Taxonomy Database (GTDB v220)^84^ representative genomes using the Infernal 1.1.5 cmsearch program^85^ with the corresponding Rfam 14.10^86^ covariance model (RF02540 for archaea and RF02541 for bacteria) using an e-value cutoff of 1e-20. For each genome, we selected the single most significant hit, additionally filtering for hits with <10% non-GC/AU/GU base pairs in regions corresponding to helices in the Rfam consensus secondary structure to minimize hits to pseudogenes. This resulted in 23S rRNA sequences for 3,686 archaeal and 58,032 bacterial species (Supplemental Files 1-2), which were aligned to the respective Rfam model using the Infernal cmalign program.

To plot PTC nucleotide frequencies in Thermoproteota, we used WebLogo3^87^ webserver, only including 23S rRNA sequences with no gaps in the shown PTC region. To calculate the Shannon entropy of PTC positions in bacteria and archaea, we only considered 23S rRNA sequences with no gaps in the plotted PTC regions (2,435 archaeal and 40,562 bacterial species). Shannon entropy was calculated from nucleotide counts at each position with a pseudocount of 1e-10 added to each sum to avoid division by zero (Supplemental File 3).

uL3 sequences were analyzed using an existing alignment^88^. The distribution of novel rProteins was established by iterative searches using the homology search tool HMMER 3.4^89^ hmmsearch program against the predicted proteomes of representative genomes for all GTDB archaeal species. The first search was performed using a profile HMM built from the *P. calidifontis* sequence, keeping only the top hit per genome. Hits across all archaea were examined manually, and a conservative set of well-aligning and full-length hits was selected within an e-value plateau. Selected sequences were aligned with mafft v7.525^90^ using default settings and profile HMMs were built using HMMER hmmbuild program to use in the subsequent search. This process was repeated until no new full-length hits were recovered. Detailed information on which sequences were included in each search can be found in Supplemental File 4 and the final homolog sequences are in Supplemental Files 5-8.

Dri homologs were identified by first using a profile HMM built from the *P. calidifontis* Dri sequence to search UniProt^91^ TREMBL (release 2024_04) using the HMMER program hmmscan. From these results, an initial Dri profile HMM was built from 22 full-length hits from Thermoproteaceae with e-values <1e-90 that were aligned using mafft with default settings. This initial Dri model was used to search the predicted proteomes of representative genomes for all GTDB archaeal species using hmmsearch with an e-value cutoff of 1e-10. Each hit was individually aligned to the *P. calidifontis* Dri sequence and evaluated on the following additional properties: percent alignment to *P. calidifontis* Dri N-terminal lobe (positions 14-303), percent alignment to *P. calidifontis* Dri C-terminal lobe (positions 324-640), charge of amino acids aligning to the *P. calidifontis* Dri N-terminal lobe ribosome binding surface (positions 43-44, 76-77, 203-210, 215, 224-225, and 227-241), charge of amino acids aligning to the *P. calidifontis* Dri C-terminal lobe P-site loop including a 3 amino acid buffer on either side (positions 544-579), and length of regions aligning to the *P. calidifontis* Dri C-terminal lobe P-site loop (positions 544-579), C-terminal lobe A-site loop (positions 394-425), and the N-terminal lobe P-site loop (positions 205-236) including a 3 amino acid buffer on either side. A final set of 92 full-length Dri homologs was selected by filtering hits on >75% N-lobe and >75% C-lobe alignment, and e-value <1e-60. All of these full-length Dri homologs had sequence features consistent with the mode of ribosome binding seen in *P. calidifontis* such as a positively charged ribosome binding region on the N-terminal lobe and the presence and positive charge of the P-site loop on the C-terminal lobe.

The 92 full-length Dri homologs were aligned using mafft with default settings, and then split into N-lobe and C-lobe only regions corresponding to the *P. calidifontis* positions 14-303 and 324-640, respectively. These alignments were used to build single-lobe profile HMMs which were used to search all GTDB archaeal species using hmmsearch. Dri N-terminal lobe-only homologs were selected by filtering based on e-value <1e-30, ≥50% N-terminal lobe and <50% C-terminal lobe alignment, P-site loop length ≥13 amino acids, ribosome binding surface net charge ≥0 and basic amino acids ≥4. E-value and lobe alignment filters alone resulted in 647 sequences with only 9 removed with the additional filters. Dri C-terminal lobe-only homologs were selected by filtering based on e-value <1e-30, ≥50% C-terminal lobe and <50% N-terminal lobe alignment, A-site loop length ≥10 amino acids, P-site loop length ≥25 amino acids, P-site loop net charge ≥0 and basic amino acids ≥4. E-value and lobe alignment alone resulted in 782 sequences with 75 removed with the additional filters. Information on the identified Dri homologs can be found in Supplemental File 9 and sequences can be found in Supplemental Files 10-12.

The distribution of Dri homologs was plotted on the GTDB v220 archaeal phylogenetic tree using the Interactive Tree of Life (iTOL v6) webserver^92^. To tabulate the frequency of single-lobed Dri gene arrangements, we only considered genomes where both Dri N-terminal and C-terminal single lobe homologs were found and additionally restricted to genomes where at least one of the homologs occurs >2000 nucleotides from the edge of a contig, resulting in 388 genomes.

### Methanosarcina acetivorans ribosome preparation

*Methanosarcina acetivorans* was grown at 37 °C without shaking in bicarbonate-buffered high salt liquid medium with 50 mM trimethylamine hydrochloride and N_2_/CO_2_ (80/20) in the headspace^93^. Cultures were maintained at stationary phase for four days and on the fourth day were placed on ice for 15 minutes. Cells were pelleted at 5,000 *xg* for 15 minutes and cell pellets were flash frozen and stored at –80 °C. The cell pellet was then thawed, resuspended in 15 mL *Ma* lysis buffer (50 mM K_x_H_y_PO_4_ pH 7, 10 mM MgCl_2_) with 20 units of RQ1 DNase (Promega) and a Pierce protease inhibitor mini tablet (ThermoFisher), and incubated at room temperature for 10 minutes. After incubation, MgCl_2_ was added to a final concentration of 20 mM and the lysate was clarified by centrifugation at 12,000 rpm (29,000 *xg*) for 45 minutes in an F14-14 x 50 cy rotor (ThermoFisher). The lysate was then loaded onto a sucrose cushion with 40 mL of *Ma* buffer B (30 mM HEPES-KOH pH 7, 60 mM KCl, 20 mM MgCl_2_, 2 mM DTT) with 1 M sucrose, and the cushion was spun in a Ti-45 rotor at 38,000 rpm (113,000 *xg*) for 16 hours. The ribosome pellet was resuspended in 500 µL *Ma* buffer B and 1 mg of ribosomes was loaded onto a 25-40% sucrose gradient in *Ma* buffer B. The sucrose gradient was placed in an SW-32 rotor (Beckman-Coulter) and centrifuged at 26,000 rpm (83,000 *xg*) for 16 hours. The sucrose gradient was then fractionated on a BioComp fractionator and the 70S peak was collected for mass spectrometry.

### Protein Mass Spectrometry

For the identification of *P. calidifontis* 50S rProteins, 5 µg of 50S subunits were buffer exchanged into 100 mM Tris-HCl, pH 8.5 with 8 M urea. TCEP was added to a concentration of 5 mM and the solution was incubated at room temperature for 20 minutes. 10 mM iodoacetamide was added to the solution, and the sample was incubated for an additional 15 minutes in the dark. The sample was then diluted with 100 mM Tris-HCl pH 8.5 until the urea concentration was 6 M. 50 units of Benzonase were added to the solution and the reaction was incubated at 35 °C for 15 minutes. The sample was then diluted again with 100 mM Tris-HCl pH 8.5 until the urea concentration was 2 M. 1 mM CaCl_2_ and 1 µg trypsin/LysC were added to the sample and the reaction was incubated overnight at 37 °C. 5% formic acid was then added to quench the digestion reaction.

To identify *P. calidifontis* 30S or *M. acetivorans* 70S rProteins, 20 µg of 30S subunits or 70S ribosomes were dried under vacuum and the pellet was resuspended with 0.1% RapiGest SF in 50 mM ammonium bicarbonate and 5 mM DTT. The solution was incubated at 80 °C for 30 minutes and then cooled to room temperature. 15 mM iodoacetamide was added and the sample was incubated in the dark for 30 minutes. The sample was then sonicated on ice using a handheld ultrasonic homogenizer with a 2 mm probe (Shengwin) at 40% power for three 30 second cycles with a 30 second rest period in-between cycles. 50 units of Benzonase were added and the sample was incubated at 35 °C for 15 minutes. The sample was then sonicated again using the above settings. 0.5 µg of chymotrypsin (*P. calidifontis* 30S) or trypsin/LysC (*M. acetivorans* 70S) was added to the sample and the reaction was incubated at 37 °C for 1 hour. The sample was sonicated a third time using the above settings, an additional 0.5 µg of chymotrypsin or trypsin/LysC was added to the sample, and the reaction was incubated at 37 °C overnight. Finally, 5% formic acid was added to quench the digestion reaction.

Mass spectrometry was performed at the Vincent J. Coates Proteomics/Mass Spectrometry Laboratory at the University of California, Berkeley. Multidimensional protein identification technique (MudPIT) was performed as described^94,95^. Briefly, a 2D nano-LC column was packed in a 100-μm inner diameter glass capillary with an integrated pulled emitter tip. The column consisted of 10 cm of ReproSil-Gold C18-1.9 μm resin (Dr. Maisch GmbH) and 4 cm of strong cation exchange resin (Partisphere, HiChrom). The column was loaded and conditioned using a pressure bomb and then coupled to an electrospray ionization source mounted on a Thermo-Fisher LTQ XL linear ion trap mass spectrometer. An Agilent 1200 HPLC equipped with a split line to deliver a flow rate of 1 µL/minute was used for chromatography. Four buffers were used to elute peptides: Buffer A (5% acetonitrile, 0.02% heptafluorobutyric acid (HFBA)), buffer B (80% acetonitrile, 0.02% HFBA), buffer C (250 mM NH_4_(OAc), 0.02% HFBA), and buffer D (500 mM NH_4_(OAc), 0.02% HFBA). Peptides were eluted in four steps, with the system equilibrated in buffer A, step 1: 0-80% buffer B in 70 minutes, step 2: 0-50% buffer C in 5 minutes and 0-45% buffer B in 100 minutes, step 3: 0-100% buffer C in 5 minutes and 0-45% buffer B in 100 minutes, and step 4: 0-100% buffer D in 5 minutes and 0-45% buffer B in 160 minutes. Collision-induced dissociation (CID) spectra were collected for each *m/z*. Protein identification, quantification, and analysis were done with PEAKS (Bioinformatics Solutions Inc.) with the following parameters: semi-specific cleavage specificity at the C-terminal site of R and K, allowing for 5 missed cleavages, precursor mass tolerance of 3 Da, and fragment ion mass tolerance of 0.6 Daltons. Methionine oxidation and phosphorylation of serine, threonine, and tyrosine were set as variable modifications and cysteine carbamidomethylation was set as a fixed modification. Peptide hits were filtered using a 1% false discovery rate (FDR).

### CRISPR-Cas9 Editing

CRISPR-Cas9 editing of *E. coli* was conducted with a modified pCas/pTargetF system^56^. sgRNA sequences were inserted in the pEcgRNA plasmid using the corresponding primer set (Supplemental Table 8) and the InFusion cloning kit. NEB Express I^q^ cells (chloramphenicol (CAM) resistant) were transformed with the pEcCas plasmid, which encodes for Cas9, the inducible λ-Red system, and kanamycin resistance (KAN). Positive transformations were grown overnight in LB with 50 µg/mL KAN and 10 µg/mL CAM (LB-KAN-CAM). 400 µL of the overnight culture was added to 40 mL of fresh LB-KAN-CAM. The culture was then grown at 37 °C with shaking until an OD_600_ of 0.4 was obtained. Expression of the λ-Red system was induced with 10 mM arabinose, and the culture was incubated for another 15 minutes at 37 °C with shaking. Cells were pelleted by centrifugation at 4,000 *xg* for 15 minutes, washed with ice-cold sterile water, pelleted again by centrifugation at 4,000 *xg* for 15 minutes, and resuspended in 400 µL of cold sterile 10% glycerol. 50 µL of cells were electroporated with 100 ng of the pEcgRNA plasmid, which contains the sgRNA (Supplemental Table 9) and spectinomycin (SPEC) resistance, and 400 ng of double stranded linear donor dsDNA (Supplemental Table 10). The cells were then selected on LB-KAN-CAM-SPEC agar plates with 100 µg/mL SPEC. PCR amplifications of the *rplC* gene were Sanger sequenced (Elim Bio) (Extended Data Fig. 3b) to identify positive clones.

To cure *E. coli* strains of the pEcCas and pEcgRNA plasmids, edited colonies were used to inoculate 2 mL of LB-KAN-CAM containing 40 mM rhamnose, and the culture was grown overnight at 37 °C with shaking. 10 µL of the overnight culture was then used to inoculate a 2 mL LB-CAM culture that was grown for 6 hours at 37 °C with shaking. The culture was diluted 1:300 in SOC media and 5 µL of the dilution was spread on an LB-CAM-agar plate with 1% sucrose and incubated at 37 °C overnight. Plasmid curing was confirmed by replica plating colonies onto LB-CAM, LB-KAN, and LB-SPEC agar plates.

For growth measurements of edited *E. coli* strains, an LB-CAM culture was grown overnight at 37 °C. The culture was then diluted to OD 0.01 in LB with no antibiotics, and 200 µL of the diluted cells were placed into a sterile 96-well plate. The OD_600_ was measured every 10 minutes using a Park Plate Reader (Tecan) at 37 °C with shaking.

### *E. coli* Ribosome Expression and Purification

A modified version of the pLK35 plasmid^96^, which contains an IPTG inducible tac promoter and *E. coli* 23S rRNA with the MS2-tag from Nissley et al.^57^ inserted in helix H98, was used to express mutant *E. coli* ribosomes. The pLK35 plasmid was linearized with primers PTC_InFusion_F/R (Supplemental Table 8), and mutations were introduced into the 23S rRNA using the InFusion cloning kit (Takara bio) and corresponding gene block (Twist) (Supplemental Table 10). Sequences were confirmed with whole plasmid sequencing (Elim Bio).

Untagged *E. coli* 30S subunits were purified from *E. coli* MRE600 as previously described^25^. MS2-tagged *E. coli* ribosomes were purified as previously described with adaptions^25^. pLK35 plasmids were transformed into NEB Express I^q^ cells. Transformants were grown in LB media overnight, and the next morning, they were diluted 1:100 in 1 L of LB media with 100 µg/mL ampicillin. The culture was grown at 37 °C with shaking until it reached an OD_600_ of 0.6. rRNA expression was induced with 0.5 mM IPTG and the cultures were grown for an additional three hours at 37 °C. Cells were pelleted and resuspended in 30 mL of *Ec* buffer A (20 mM Tris-HCl pH 7.5, 100 mM NH_4_Cl, 10 mM MgCl_2_) with a Pierce protease inhibitor tablet (ThermoFisher). Cells were lysed by sonication and the lysate was clarified by centrifugation at 12,000 rpm (29,000 *xg*) for 45 minutes in an F14-14 x 50 cy rotor (ThermoFisher). The lysate was then loaded onto a sucrose cushion with 24 mL of *Ec* buffer B (20 mM Tris-HCl pH 7.5, 500 mM NH_4_Cl, 10 mM MgCl_2_) with 0.5 M sucrose and 17 mL of *Ec* buffer C (20 mM Tris-HCl pH 7.5, 60 mM NH_4_Cl, 6 mM MgCl_2_) with 0.7 M sucrose. Ribosomes were pelleted by centrifugation at 27,000 rpm (57,000 *xg*) for 16 h at 4 °C in a Ti-45 rotor (Beckman-Coulter). Ribosomes pellets were then resuspended in dissociation buffer (20 mM Tris-HCl pH 7.5, 60 mM NH_4_Cl, 1 mM MgCl_2_).

10 mg of MBP-MS2 fusion protein, purified as previously described^69^, was loaded onto a 5 mL MBP Trap column (Cytiva) that was pre-equilibrated with MS2-150 buffer (20 mM HEPES pH 7.5, 150 mM KCl, 1mM EDTA, 2 mM 2-mecaptoethanol). The column was washed with 5 CV of *Ec* buffer A-1 (20 mM Tris-HCl pH 7.5, 100 mM NH_4_Cl, 1 mM MgCl_2_), and the resuspended ribosomes were loaded onto the column. The column was then washed with 5 CV of *Ec* buffer A-1 and 10 CV of *Ec* buffer A-250 (20 mM Tris-HCl pH 7.5, 250 mM NH_4_Cl, 1 mM MgCl_2_). Tagged *E. coli* 50S subunits were eluted with 10 mL of *Ec* elution buffer (20 mM Tris-HCl pH 7.5, 100 mM NH_4_Cl, 1 mM MgCl_2_, 10 mM maltose). The sample was then concentrated in a 100 kDa cut-off spin filter (Millipore) and washed with *Ec* buffer A-1. 50S subunits were quantified using the approximation of 1 A_260_ = 36 nM, flash frozen, and stored at –80 °C.

Endogenous *E. coli* 50S subunit contamination was quantified using semi-quantitative RT-PCR. 50 pmol of MS2-purified 50S subunits were denatured at 95 °C for 5 minutes, and the rRNA was precipitated with 4M LiCl. 75 ng of rRNA was reverse transcribed and amplified with 8 PCR cycles using the OneStep RT-PCR kit (Qiagen) and primers MS2_quant_F and MS2_quant_R (Supplemental Table 8). DNA products were resolved on a 10% TBE gel, visualized with SYBR gold stain (Thermo Fisher), and quantified using Image J software^97^.

### *E. coli* Ribosome Activity Assays

HiBiT^65^ *in vitro* translation assays were conducted as previously described^25^. Briefly, an *in vitro* translation reaction was assembled with the ΔRibosome PURExpress kit (NEB): 3.2 µL solution A (NEB), 1 µL factor mix (NEB), 250 nM MS2-tagged 50S ribosomal subunit, 500 nM untagged 30S ribosomal subunit, 1 U/µL murine RNAse inhibitor (NEB), 400 nM 11S NanoLuc protein (purified as described in^25^), 1:50 (v/v) dilution of Nano-Glo substrate (Promega), and 1 ng/µL of DNA template encoding the HiBiT peptide (Supplemental Table 10). 2 µL of the mixture was then placed in a well of a 384-well plate, and luminescence was measured in a Spark plate reader (Tecan) at 37 °C incubation. Ribosome activity was determined as the slope of the initial linear increase in the signal of each *in vitro* translation reaction.

## Data availability

Ribosome coordinates have been deposited in the Protein Data Bank for the *P. calidifontis* 50S-Dri complex (9E6Q), 70S consensus reconstruction (9E71), and 70S-Dri complex (9E7F). Cryo-EM maps have been deposited in the Electron Microscopy Data Bank for the 50S-Dri complex (EMD-47578), 70S consensus reconstruction (EMD-47604, EMD-47605, EMD-47606, EMD-47611, EMD-47617, EMD-47628), and 70S-Dri complex (EMD-47662, EMD-47664, EMD-47666, EMD-47667, EMD-47668).

## Contributions

A.J.N., P.I.P., and J.H.D.C. conceptualized the project. A.J.N and B.E.D. cultured archaea. A.J.N. acquired cryo-EM data and conducted image analysis and modeling. A.J.N. and R.W.K. performed biochemical experiments. Y.S. performed bioinformatic analysis. A.J.N. and Y.S. prepared figures. A.J.N. wrote the initial draft and all authors reviewed and edited the manuscript. J.H.D.C., J.F.B., and D.D.N were responsible for funding and supervised the project.

## Ethics declarations

J.H.D.C. is a founder and board and SAB member of Initial Therapeutics. The remaining authors declare no competing interests.

## Acknowledgements

We thank Dan Toso, Ravindra Thakkar, and Paul Tobias for assistance with cryo-EM data collection (Cal-Cryo). This work was funded by the NSF Center for Genetically Encoded Materials (C-GEM) (CHE-2002182). Y.S. is a Don Brown Awardee of the Life Sciences Research Foundation. D.D.N. is a Chan Zuckerberg Biohub – San Francisco Investigator. This work used the Vincent J. Coates Proteomics/Mass Spectrometry Laboratory Core Facility, RRID:SCR_025852.

---

**Extended Data Fig. 1.**
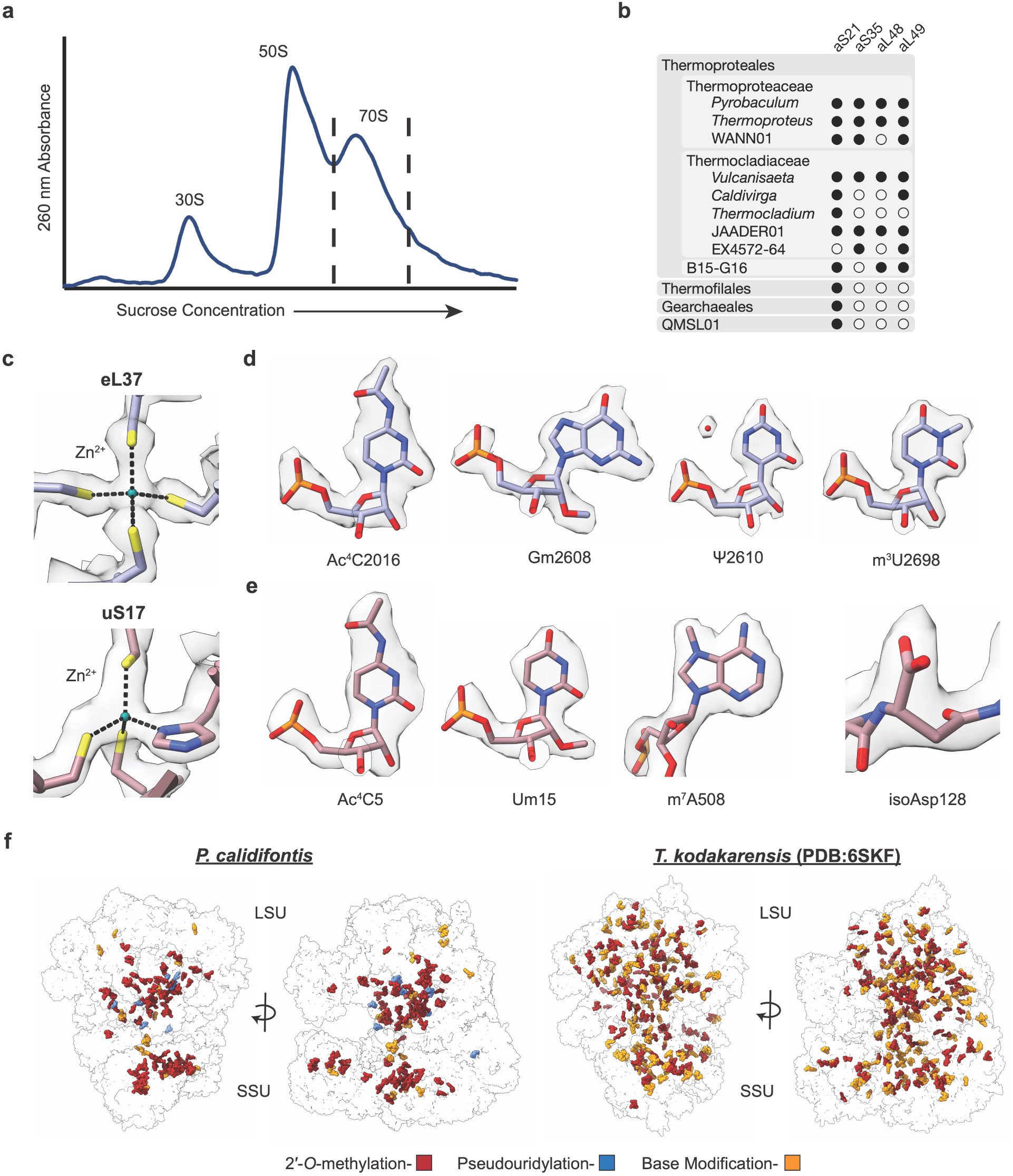
Cryo-EM sample preparation and analysis of post-transcriptional modifications in *P. calidifontis*. **a**, Sucrose gradient fractionation of *P. calidifontis* ribosomes. Dashed lines bracket the sample that was collected for cryo-EM analysis. **b**, Distribution of archaea-specific rProteins across four orders and further subclades in the class Thermoprotei. Closed and open circles represent clades where rProtein homologs were or were not identified, respectively. **c**, Zinc binding sites in LSU rProtein eL37 and SSU rProtein uS17 with cryo-EM density. **d-e,** Cryo-EM density for representative post-transcriptional and post-translational modifications in the LSU (**d**) and SSU (**e**). **f**, Distribution of post-transcriptional modifications in *P. calidifontis* (left) and *T. kodakarensis* PDB:6SKF^22^ (right). Note that pseudouridines are not modeled in *T. kodakarensis*.

**Extended Data Fig. 2.**
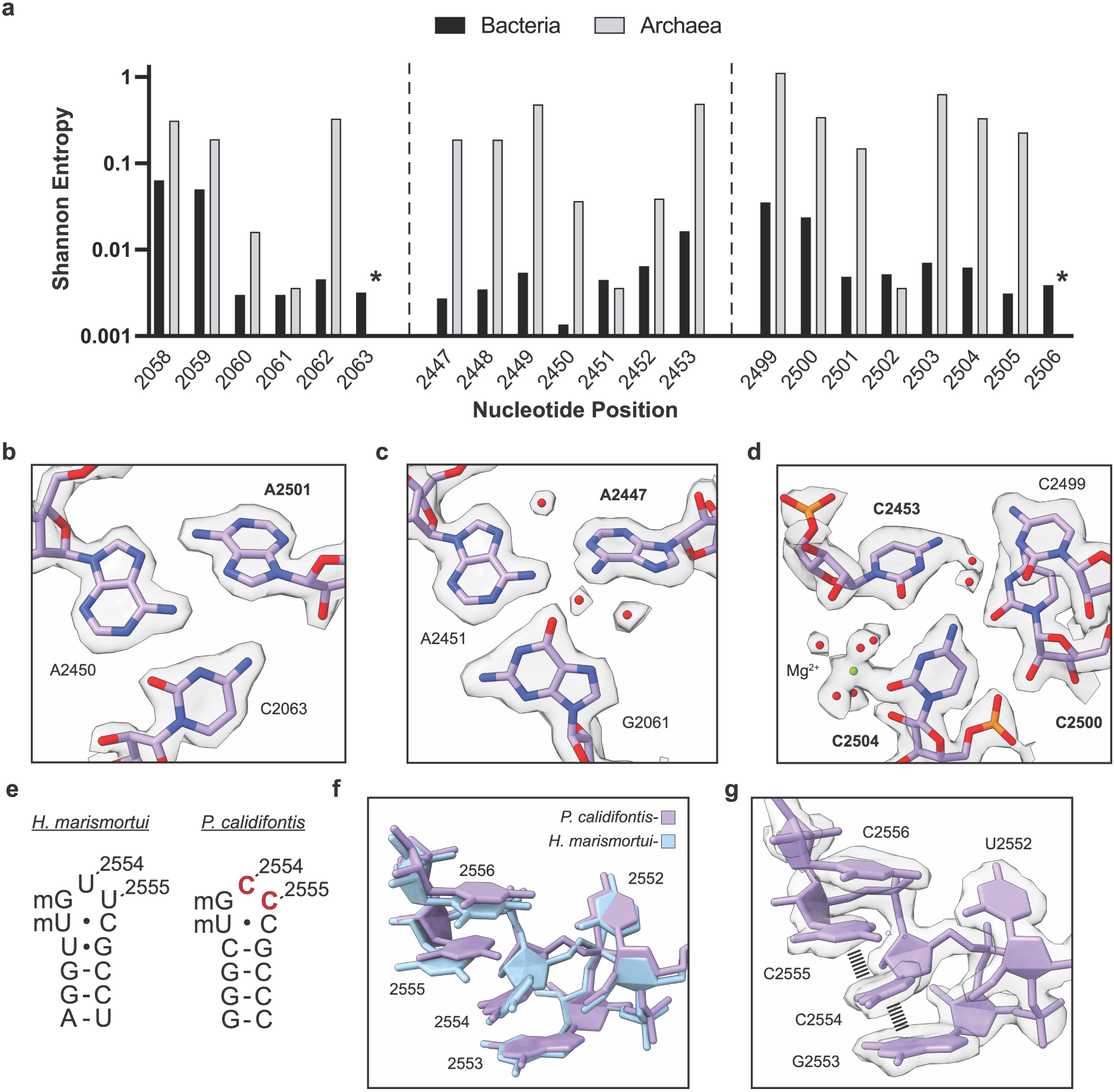
Sequence and structural diversity in the *P. calidifontis* PTC. **a**, Shannon entropy plot for PTC nucleotides in GTDB bacteria and archaea. Positions indicated by a * have a Shannon entropy value of zero for archaea. A Shannon entropy value of zero corresponds to complete conservation of the nucleotide, while a value of 1.386 corresponds to an equal distribution of all four nucleotides. **b-d**, Cryo-EM density for regions of the PTC shown in Figure 2. Water molecules and magnesium ions are shown as red and green spheres respectively. **e**, Secondary structures of the *H. marismortui* and *P. calidifontis* A loops. Positions with rare sequence variation, 2554 and 2555, are highlighted in red in the *P. calidifontis* A loop. **f**, Comparison of the apical region of the A loop in *H. marismortui* PDB:1S72^46^ (blue) and *P. calidifontis* (purple). **g**, Cryo-EM density for the *P. calidifontis* A loop. Base stacking interactions are shown as dashed lines.

**Extended Data Fig. 3.**
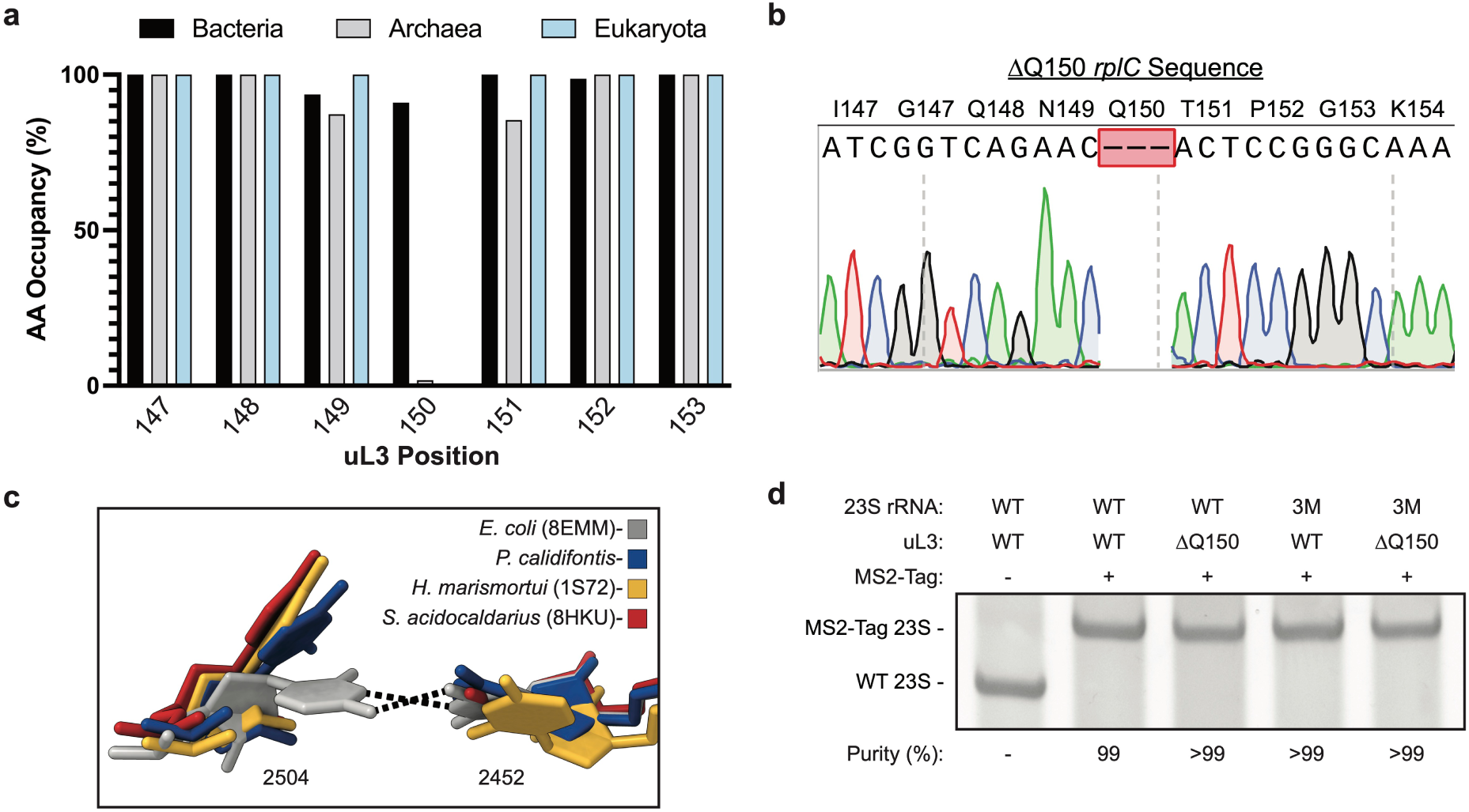
Differences in uL3 and the PTC between the domains of life. **a**, Amino acid occupancy in an alignment of uL3 sequences from representative species in bacteria, archaea, and eukaryotes^88^. **b**, Sequencing chromatogram for the *rplC* gene from ΔQ150-uL3 *E. coli*, showing the deletion of residue Q150. The chromatogram trace was broken at the site of deletion for clarity. **c**, Comparison of the positions of 23S rRNA nucleotides 2452 and 2504 in *E. coli* and archaeal ribosomes. **d**, RT-PCR purity assay for MS2-tagged *E. coli* ribosomes used in this study. Sample purity was quantified using the band intensities of the upper (MS2-tagged ribosome) and lower (WT ribosome) bands.

**Extended Data Fig. 4.**
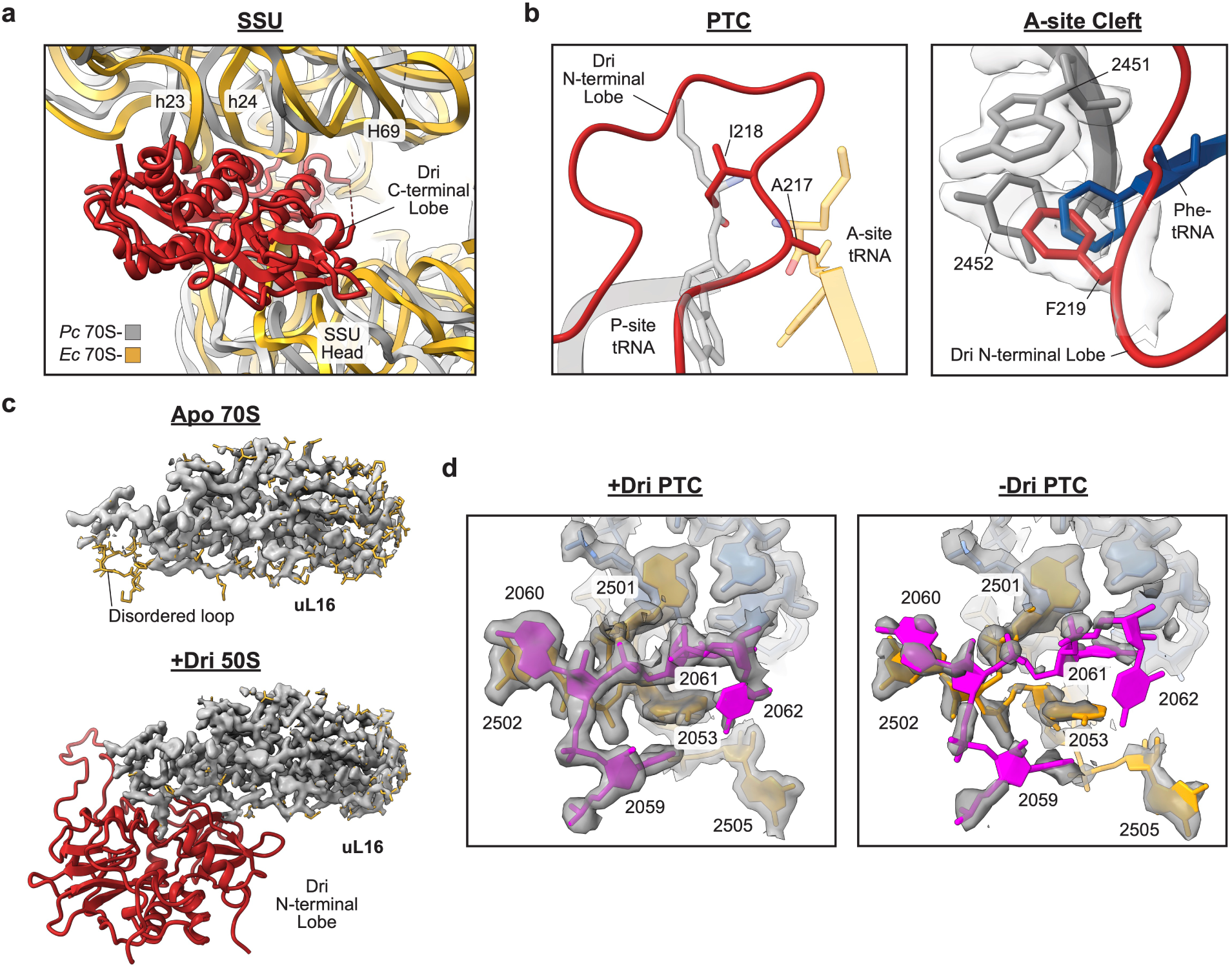
Dri interactions with the ribosome. **a**, Dri (red) binds to the rotated state of the *P. calidifontis* ribosome (grey). An *E. coli* ribosome in the unrotated state (PDB:8EMM^61^) is overlayed for reference (grey). **b**, (left) Dri residues occupy the positions of the A and P-site amino acids in the PTC. The A and P-site tRNAs from PDB:8EMM^61^ are overlayed on the *P. calidifontis* ribosome (right). Dri residue F219 occupies the A-site cleft in a position similar to phenylalanyl-tRNA^Phe^ (PDB: 1VY4^62^). **c**, In the absence of the Dri protein, a loop of rProtein uL16 is disordered (top). Cryo-EM density is shown in gray. Upon binding of the Dri N-terminal lobe to the LSU, the loop of uL16 becomes ordered (bottom). **d**, Cryo-EM density for PTC nucleotides in the 50S-Dri complex (left) or the composite 70S complex (right).

**Extended Data Fig. 5.**
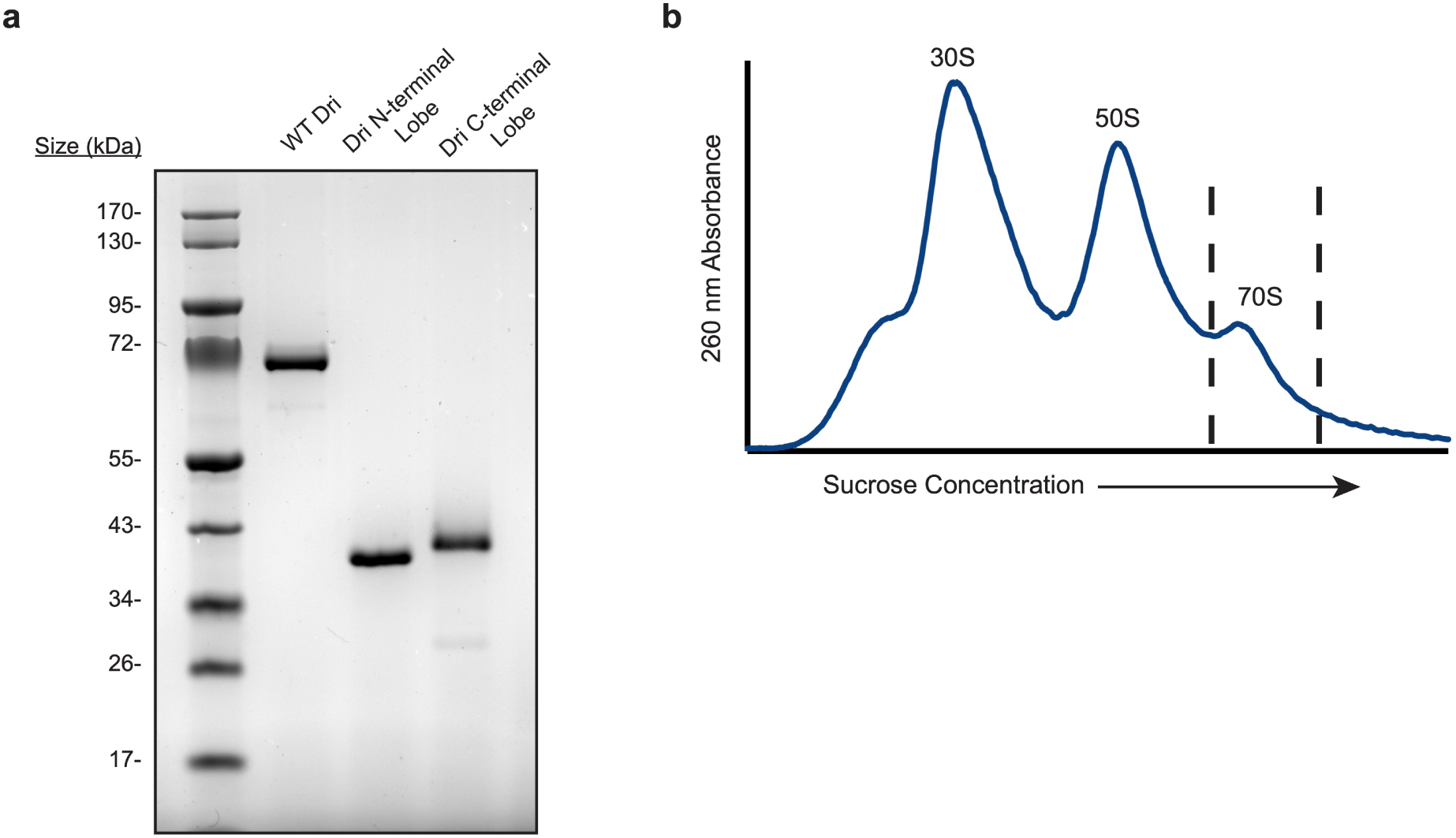
Recombinant Dri Expression and Purification of *M. acetivorans* ribosomes. **a**, Purified Dri proteins resolved on a 4-12% polyacrylamide gel stained with Coomassie brilliant blue. **b**, Sucrose gradient fractionation of *M. acetivorans* ribosomes. Dashed lines bracket the sample that was collected for mass spectrometry.

**Extended Data Table 1.** Dri homologs in archaeal phyla.

| Phylum | # species | Both Lobes | N-lobe | C-lobe |
| --- | --- | --- | --- | --- |
| Aenigmataarchaeota | 165 | 0 | 0 | 1 |
| Altiarchaeota | 33 | 0 | 17 | 0 |
| Asgardarchaeota | 296 | 1 | 1 | 5 |
| B1Sed10-29 | 19 | 0 | 19 | 10 |
| EX4484-52 | 34 | 0 | 0 | 0 |
| Hadarchaeota | 27 | 0 | 0 | 15 |
| Halobacteriota | 1223 | 0 | 343 | 402 |
| Huberarchaeota | 1 | 0 | 0 | 0 |
| Hydrothermarchaeota | 23 | 0 | 14 | 15 |
| Iainarchaeota | 101 | 0 | 0 | 0 |
| JACRDV01 | 1 | 0 | 1 | 1 |
| Methanobacteriota | 244 | 0 | 81 | 82 |
| Methanobacteriota_A | 35 | 0 | 0 | 0 |
| Methanobacteriota_B | 116 | 0 | 2 | 0 |
| Micrarchaeota | 331 | 0 | 0 | 0 |
| Nanoarchaeota | 847 | 0 | 0 | 0 |
| Nanohaloarchaeota | 29 | 0 | 0 | 0 |
| SpSt-1190 | 10 | 0 | 0 | 0 |
| Thermoplasmatota | 1064 | 0 | 1 | 0 |
| Thermoproteota | 1264 | 90 | 151 | 173 |
| Undinarchaeota | 6 | 0 | 2 | 0 |
| Total | 5869 | 91 | 632 | 704 |

