## Supplemental Information for "Structure of an Archaeal Ribosome with a Divergent Active Site"

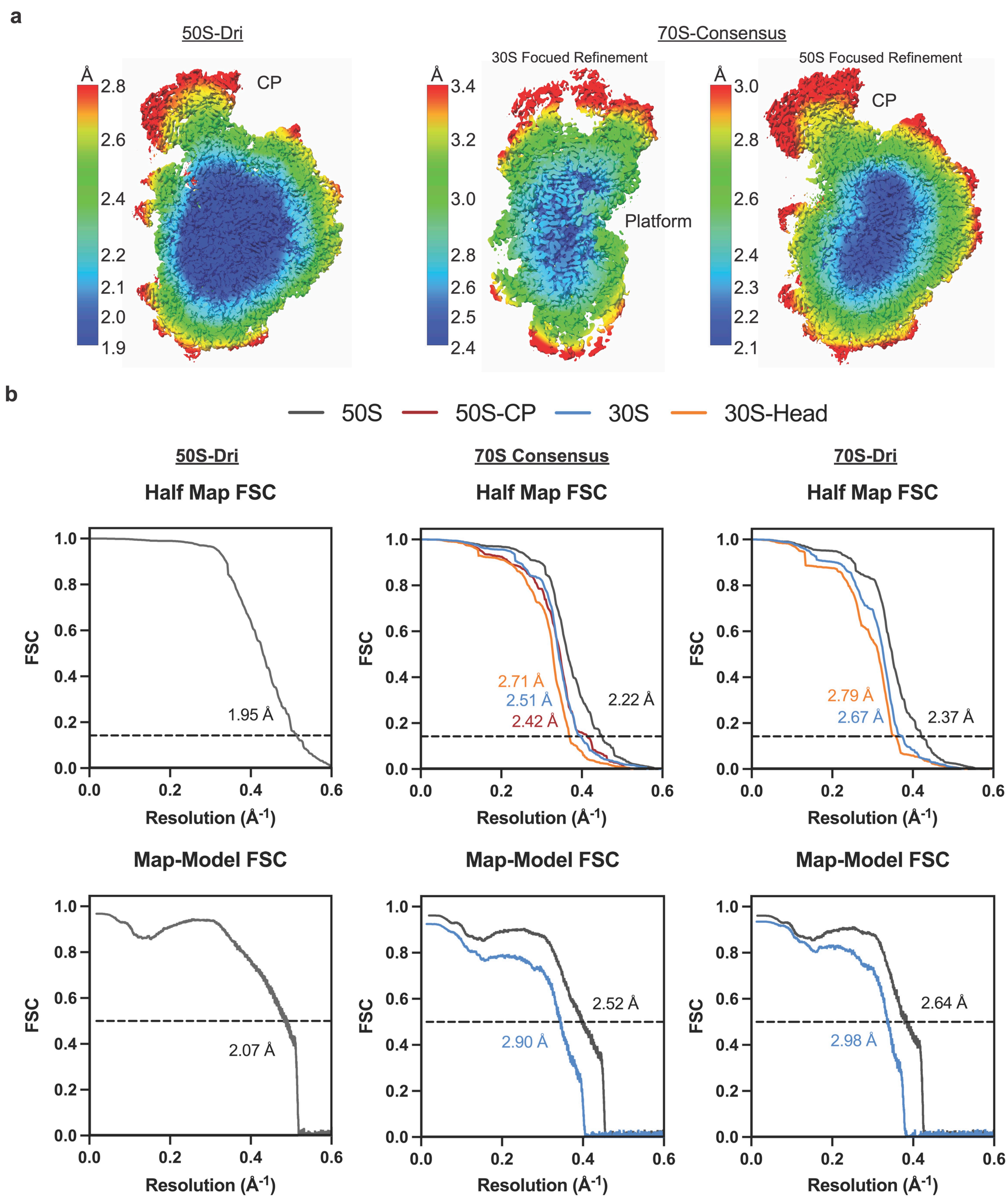


**Supplemental Fig. 1** Resolution of cryo-EM maps. **a**, Local resolution plotted on the 50S (left) or 70S consensus (right) reconstructions. The 30S platform and 50S central protuberance (CP) are labelled. **b**, Half map (top) and map to model (bottom) FSCs for cryo-EM maps and models in this study.

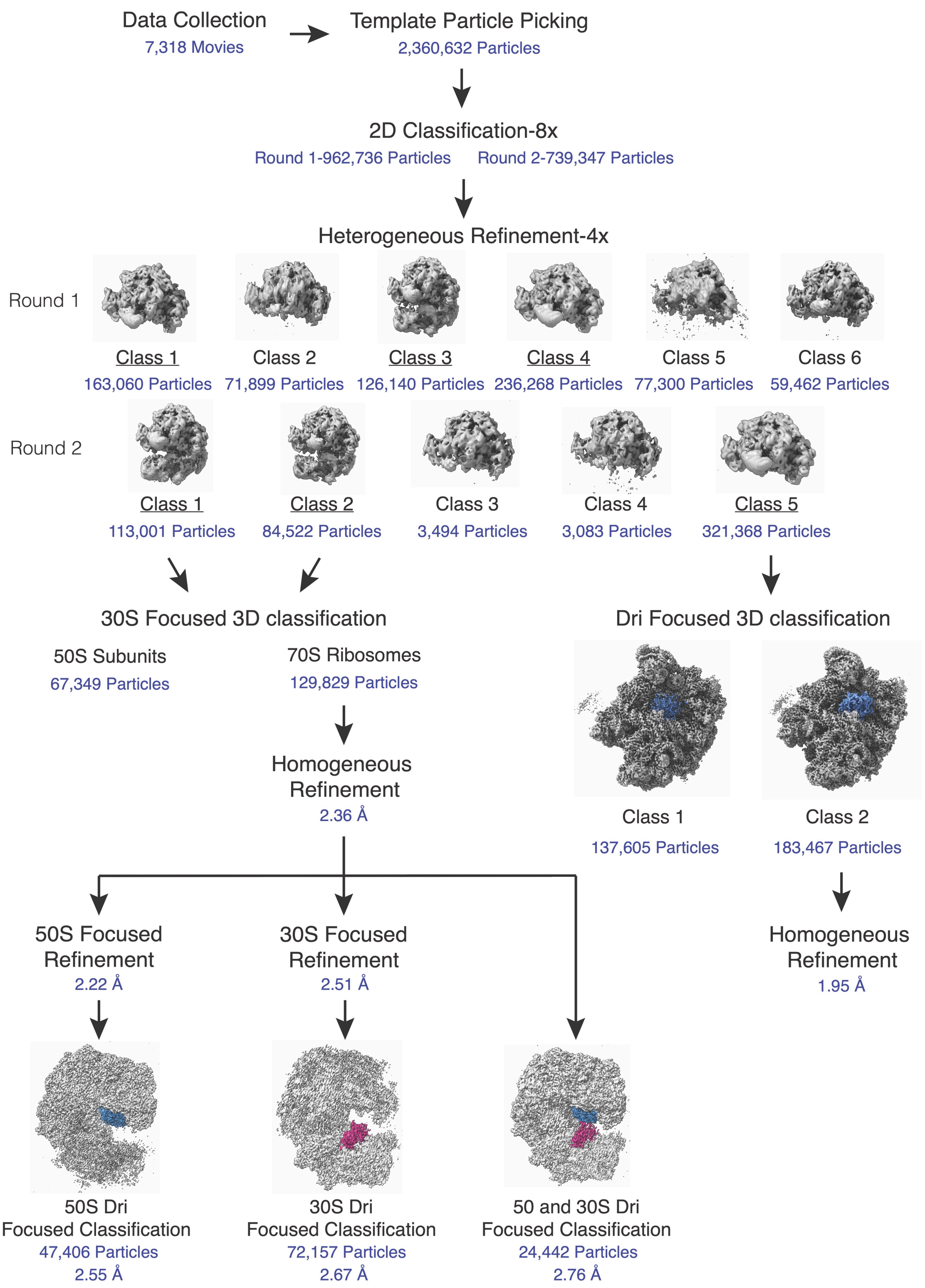


**Supplemental Fig. 2** Cryo-EM processing workflow.

**Supplemental Table 1.** ***P. calidifontis*** **Large Ribosomal Subunit Proteins Identified by Mass Spectrometry**

| Accession | -10LgP | Coverage (%) | #Peptides | #Unique Peptides | Average Mass | Description |
|---|---|---|---|---|---|---|
| A3MTT0\|A3MTT0_PYRCJ | 359.93622 | 76.85 | 20 | 20 | 11174.706 | Large ribosomal subunit protein P1 |
| A3MX83\|RL1_PYRCJ | 243.00838 | 32.88 | 7 | 7 | 25074.805 | Large ribosomal subunit protein uL1 |
| A3MS41\|RL2_PYRCJ | 249.16068 | 22.95 | 7 | 7 | 26193.252 | Large ribosomal subunit protein uL2 |
| A3MWI4\|RL3_PYRCJ | 375.15442 | 39.05 | 18 | 18 | 37181.547 | Large ribosomal subunit protein uL3 |
| A3MWI5\|A3MWI5_PYRCJ | 384.69888 | 56.14 | 21 | 21 | 31547.998 | Large ribosomal subunit protein uL4 |
| A3MXP8\|A3MXP8_PYRCJ | 291.17993 | 38.76 | 9 | 9 | 20116.807 | Large ribosomal subunit protein uL5 |
| A3MWV5\|A3MWV5_PYRCJ | 421.6866 | 81.12 | 28 | 28 | 21630.463 | Large ribosomal subunit protein uL6 |
| A3MTA9\|RL7A_PYRCJ | 192.32135 | 28.86 | 4 | 4 | 15990.72 | Large ribosomal subunit protein eL8 |
| A3MX84\|A3MX84_PYRCJ | 369.66794 | 45.64 | 18 | 18 | 37480.96 | Large ribosomal subunit protein uL10 |
| A3MX82\|A3MX82_PYRCJ | 140.1296 | 18.07 | 3 | 3 | 18383.375 | Large ribosomal subunit protein uL11 |
| A3MXZ7\|A3MXZ7_PYRCJ | 419.68018 | 72.58 | 27 | 27 | 21694.902 | Large ribosomal subunit protein uL13 |
| A3MTP8\|A3MTP8_PYRCJ | 317.1464 | 48.41 | 12 | 12 | 17897.012 | Large ribosomal subunit protein eL13 |
| A3MX34\|RL14_PYRCJ | 373.60333 | 72.22 | 20 | 20 | 15742.426 | Large ribosomal subunit protein uL14 |
| A3MS76\|RL14E_PYRCJ | 203.08841 | 30.1 | 4 | 4 | 11448.51 | Large ribosomal subunit protein eL14 |
| A3MUZ1\|RL15_PYRCJ | 303.03702 | 51.28 | 10 | 10 | 17487.32 | Large ribosomal subunit protein uL15 |
| A3MWR6\|A3MWR6_PYRCJ | 252.2958 | 23.28 | 6 | 6 | 22574.615 | 50S ribosomal protein L15e |
| A3MXP3\|RL10E_PYRCJ | 384.6988 | 53.37 | 22 | 22 | 19859.158 | Large ribosomal subunit protein uL16 |
| A3MSJ3\|RL18_PYRCJ | 168.60799 | 11.71 | 3 | 3 | 23374.21 | Large ribosomal subunit protein uL18 |
| A3MXZ6\|A3MXZ6_PYRCJ | 241.26091 | 40.16 | 5 | 5 | 13201.696 | Large ribosomal subunit protein eL18 |
| A3MSJ4\|A3MSJ4_PYRCJ | 345.15067 | 40.14 | 14 | 14 | 17373.215 | Large ribosomal subunit protein eL19 |
| A3MX77\|RL18A_PYRCJ | 313.1881 | 84.62 | 10 | 10 | 8913.517 | Large ribosomal subunit protein eL20 |
| A3MWX9\|RL21_PYRCJ | 263.81876 | 46.46 | 6 | 6 | 11294.232 | Large ribosomal subunit protein eL21 |
| A3MTL6\|RL22_PYRCJ | 314.46808 | 39.67 | 11 | 11 | 21268.16 | Large ribosomal subunit protein uL22 |
| A3MWI6\|A3MWI6_PYRCJ | 162.22151 | 25.93 | 2 | 2 | 9290.063 | Large ribosomal subunit protein uL23 |
| A3MWZ8\|RL24_PYRCJ | 294.34494 | 42.97 | 9 | 9 | 14555.221 | Large ribosomal subunit protein uL24 |
| A3MTA8\|RL24E_PYRCJ | 298.51492 | 64.52 | 9 | 9 | 7085.4673 | Large ribosomal subunit protein eL24 |
| A3MTL2\|A3MTL2_PYRCJ | 324.05475 | 58.23 | 12 | 12 | 9136.798 | Large ribosomal subunit protein uL29 |
| A3MU87\|RL30_PYRCJ | 113.587166 | 12.85 | 2 | 2 | 20258.785 | Large ribosomal subunit protein uL30 |
| A3MXZ2\|RL30E_PYRCJ | 245.51628 | 63.37 | 6 | 6 | 10871.856 | Large ribosomal subunit protein eL30 |
| A3MX75\|RL31_PYRCJ | 286.8368 | 49.45 | 8 | 8 | 10643.573 | Large ribosomal subunit protein eL31 |
| A3MSJ5\|A3MSJ5_PYRCJ | 321.42322 | 34.64 | 12 | 12 | 18131.469 | Large ribosomal subunit protein eL32 |
| A3MV68\|RL34_PYRCJ | 251.49355 | 33.33 | 5 | 5 | 9350.048 | Large ribosomal subunit protein eL34 |
| A3MY38\|A3MY38_PYRCJ | 213.66008 | 48.08 | 4 | 4 | 6156.181 | Large ribosomal subunit protein eL37 |
| A3MTT1\|A3MTT1_PYRCJ | 317.47028 | 92.54 | 12 | 12 | 7939.423 | LSU ribosomal protein L38E |
| A3MUU4\|RL39_PYRCJ | 145.79361 | 35.29 | 2 | 2 | 6012.228 | Large ribosomal subunit protein eL39 |
| A3MX99\|RL40_PYRCJ | 248.84946 | 64.15 | 8 | 8 | 6339.579 | Large ribosomal subunit protein eL40 |

| | | | | | | |
|---|---|---|---|---|---|---|
| A3MUV1\|A3MUV1_PYRCJ | 278.55612 | 67.47 | 8 | 8 | 9646.363 | Large ribosomal subunit protein eL42 |
| A3MUM7\|A3MUM7_PYRCJ | 403.29984 | 79.41 | 24 | 24 | 11362.385 | Large ribosomal subunit protein eL43 |
| A3MTM7\|A3MTM7_PYRCJ | 286.37665 | 39.67 | 8 | 8 | 20927.469 | DJ-1/PfpI domain-containing protein (**aL48**) |
| A3MWU0\|A3MWU0_PYRCJ | 307.95416 | 49.46 | 11 | 11 | 10917.768 | PaREP1 domain containing protein (**aL49**) |
| A3MX58\|A3MX58_PYRCJ | 223.53175 | 6.56 | 5 | 5 | 72024.73 | Putative signal-transduction protein with CBS domains (**Dri**) |

**Supplemental Table 2. *P. calidifontis* Small Ribosomal Subunit Proteins Identified by Mass Spectrometry**

| Accession | -10LgP | Coverage (%) | #Peptides | #Unique Peptides | Average Mass | Description |
|---|---|---|---|---|---|---|
| A3MVE0\|RS3A_PYRCJ | 340.4799 | 65.32 | 21 | 21 | 24696.094 | Small ribosomal subunit protein eS1 |
| A3MS50\|RS2_PYRCJ | 434.48245 | 72.6 | 47 | 47 | 23798.812 | Small ribosomal subunit protein uS2 |
| A3MTL3\|RS3_PYRCJ | 412.854 | 73.61 | 37 | 36 | 24644.172 | Small ribosomal subunit protein uS3 |
| A3MUS9\|RS4_PYRCJ | 369.88324 | 65.41 | 23 | 23 | 18479.123 | Small ribosomal subunit protein uS4 |
| A3MWZ7\|RS4E_PYRCJ | 352.1609 | 53.16 | 19 | 19 | 26732.367 | Small ribosomal subunit protein eS4 |
| A3MU88\|RS5_PYRCJ | 378.48062 | 66.34 | 25 | 25 | 22123.717 | Small ribosomal subunit protein uS5 |
| A3MUD0\|RS6E_PYRCJ | 291.49942 | 51.66 | 13 | 13 | 16221.998 | Small ribosomal subunit protein eS6 |
| A3MS29\|A3MS29_PYRCJ | 250.44583 | 26.01 | 7 | 7 | 25475.637 | Small ribosomal subunit protein uS7 |
| A3MSJ6\|RS8_PYRCJ | 296.6819 | 49.23 | 12 | 11 | 14817.512 | Small ribosomal subunit protein uS8 |
| A3MWY2\|RS8E_PYRCJ | 251.13277 | 31.3 | 8 | 8 | 14415.8545 | Small ribosomal subunit protein eS8 |
| A3MXZ8\|A3MXZ8_PYRCJ | 183.2937 | 19.72 | 3 | 3 | 16117.1 | Small ribosomal subunit protein uS9 |
| A3MT22\|RS10_PYRCJ | 356.50195 | 62.26 | 20 | 20 | 12198.474 | Small ribosomal subunit protein uS10 |
| A3MX63\|RS11_PYRCJ | 263.05136 | 27.66 | 7 | 7 | 14963.25 | Small ribosomal subunit protein uS11 |
| A3MXZ4\|RS12_PYRCJ | 353.57635 | 80.95 | 22 | 22 | 16444.416 | Small ribosomal subunit protein uS12 |
| A3MUT0\|RS13_PYRCJ | 307.90744 | 62.09 | 15 | 15 | 16726.564 | Small ribosomal subunit protein uS13 |
| A3MSJ7\|A3MSJ7_PYRCJ | 273.35706 | 53.7 | 9 | 9 | 6414.726 | Small ribosomal subunit protein uS14 |
| A3MVD1\|RS15_PYRCJ | 200.31052 | 27.81 | 7 | 7 | 17551.531 | Small ribosomal subunit protein uS15 |
| A3MTT7\|A3MTT7_PYRCJ | 292.66876 | 45.58 | 10 | 10 | 16951.176 | Small ribosomal subunit protein uS17 |
| A3MS42\|RS17E_PYRCJ | 230.48767 | 57.75 | 6 | 6 | 8064.4707 | Small ribosomal subunit protein eS17 |
| A3MTT6\|RS19_PYRCJ | 241.82663 | 20.25 | 6 | 6 | 18147.479 | Small ribosomal subunit protein uS19 |
| A3MUU6\|A3MUU6_PYRCJ | 152.12274 | 17.09 | 2 | 2 | 17737.877 | Small ribosomal subunit protein eS19 |
| A3MSX4\|A3MSX4_PYRCJ | 261.62076 | 35.94 | 9 | 9 | 14820.318 | Small ribosomal subunit protein eS24 |
| A3MV37\|A3MV37_PYRCJ | 323.74094 | 68.18 | 17 | 17 | 12449.666 | SSU ribosomal protein S25E |
| A3MV49\|A3MV49_PYRCJ | 323.6989 | 56 | 14 | 14 | 11546.789 | SSU ribosomal protein S26E |
| A3MUV2\|RS27_PYRCJ | 178.86865 | 38.81 | 5 | 5 | 7434.8574 | Small ribosomal subunit protein eS27 |
| A3MSW3\|A3MSW3_PYRCJ | 79.18172 | 16.67 | 1 | 1 | 6098.0933 | SSU ribosomal protein S30E |
| A3MSX5\|RS27A_PYRCJ | 239.32335 | 52.31 | 7 | 7 | 7573.985 | Small ribosomal subunit protein eS31 |
| A3MY11\|A3MY11_PYRCJ | 237.6753 | 63.08 | 8 | 8 | 6947.1035 | Small zinc finger protein HVO-2753-like zinc-binding pocket domain-containing protein (**aS21**) |
| A3MX58\|A3MX58_PYRCJ | 307.99796 | 20.92 | 14 | 14 | 72024.73 | Putative signal-transduction protein with CBS domains (**Dri**) |
| WP_226952012.1 | 146.96658 | 33.82 | 2 | 2 | 7837.344 | Hypothetical protein (NCBI:WP_226952012.1) (**aS35**) |

**Supplemental Table 3. Nucleotide Numbering Conversions**

| *E. coli* Numbering | *P. calidifontis* Numbering |
|---|---|
| 2058 | 2178 |
| 2059 | 2179 |
| 2060 | 2180 |
| 2061 | 2181 |
| 2062 | 2182 |
| 2447 | 2561 |
| 2448 | 2562 |
| 2449 | 2563 |
| 2450 | 2564 |
| 2451 | 2565 |
| 2452 | 2566 |
| 2453 | 2567 |
| 2500 | 2614 |
| 2501 | 2615 |
| 2502 | 2616 |
| 2503 | 2617 |
| 2504 | 2618 |
| 2505 | 2619 |
| 2506 | 2620 |
| 2552 | 2666 |
| 2553 | 2667 |
| 2554 | 2668 |
| 2555 | 2669 |
| 2556 | 2670 |
| 2584 | 2698 |
| 2585 | 2699 |
| 2586 | 2700 |
| 2602 | 2716 |

### Supplemental Table 4. *M. acetivorans* Dri Homologs Identified by Mass Spectrometry

| Accession | -10LgP | Coverage (%) | #Peptides | #Unique Peptides | Average Mass | Description |
|---|---|---|---|---|---|---|
| Q8TH72\|Q8TH72_METAC | 36.662453 | 4.63 | 1 | 1 | 31237.389 | Uncharacterized protein (**Dri C-terminal lobe homolog**) |
| Q8TH73\|Q8TH73_METAC | 57.211075 | 6.61 | 2 | 2 | 36574.367 | Uncharacterized protein (**CBS-domain containing protein**) |

**Supplemental Table 5. Protein Sequences used in this study**

| **Name** | **Sequence** |
|---|---|
| WT Dri | MLTREELEKLAEKRVPQFVRDVVEHPPFRVTVNTTLETLAAYLRKFPVDVVPVFKSVFS<br>DEVAGVVYPHTALLLKSKKLDAKVGEFLNQPLVVKESWRIENVAEMLISESKWGAVVV<br>DEEGKFVGVVSLRGLLSALLLREPKAKSVAAVYTSIDEKKPRVGFVKAIEKVSKIFHKLV<br>GGEVDGYVVLNREGGAAGILTVWNFLKSRRWFRGSGEPRAIFGTRVTRGESKPRGVA<br>RVWRIMSRGVAVANPDTPITDVARYMATFGIYVVPVVDRNGKVIGAVTAWDVLHAYLY<br>GPKEGREDVEVKTAVEVPVSKPSMEAVIRLKPVKHVTGLRARDVMLSDVPMVNVRDPL<br>SSIRKAFLRSKSGILAVVDDSGKVVGFITRRDFMSYIAEKSLGYWRRQKGKLLVLREEAL<br>PGEQAKLAVEEGTAGDLAKSEYPTVSESATLEEVAYKMLAAGTDYVVVIDSAGQPIGVV<br>TKDELVKAYKERGRSVKVGELMTPADVAAVNPFSSLMSTTRKINAFELDGVVVVEGGNI<br>MGVVTVDDLALRPVEETLRGERVVYFTKSGVLRKATTGLSRLRYSKVGTLTALDVSRP<br>VDHVANVNADVKEVLDKLVEQGVVPVVDEQGRLVGVLNKMDIVKDLARVYVTYAMPE<br>KLAEVKQVEERAHHHHHH |
| Dri N-terminal Lobe | MLTREELEKLAEKRVPQFVRDVVEHPPFRVTVNTTLETLAAYLRKFPVDVVPVFKSVFS<br>DEVAGVVYPHTALLLKSKKLDAKVGEFLNQPLVVKESWRIENVAEMLISESKWGAVVV<br>DEEGKFVGVVSLRGLLSALLLREPKAKSVAAVYTSIDEKKPRVGFVKAIEKVSKIFHKLV<br>GGEVDGYVVLNREGGAAGILTVWNFLKSRRWFRGSGEPRAIFGTRVTRGESKPRGVA<br>RVWRIMSRGVAVANPDTPITDVARYMATFGIYVVPVVDRNGKVIGAVTAWDVLHAYLY<br>GPKEGREDVEVKTAVEVPHHHHHH |
| Dri C-terminal Lobe | MGSSHHHHHHAVIRLKPVKHVTGLRARDVMLSDVPMVNVRDPLSSIRKAFLRSKSGILA<br>VVDDSGKVVGFITRRDFMSYIAEKSLGYWRRQKGKLLVLREEALPGEQAKLAVEEGTA<br>GDLAKSEYPTVSESATLEEVAYKMLAAGTDYVVVIDSAGQPIGVVTKDELVKAYKERGR<br>SVKVGELMTPADVAAVNPFSSLMSTTRKINAFELDGVVVVEGGNIMGVVTVDDLALRP<br>VEETLRGERVVYFTKSGVLRKATTGLSRLRYSKVGTLTALDVSRPVDHVANVNADVKE<br>VLDKLVEQGVVPVVDEQGRLVGVLNKMDIVKDLARVYVTYAMPEKLAEVKQVEERA |

**Supplemental Table 6. Cryo-EM Data Collection and Processing**

| **Reconstruction** | **50S-CBS** | **70S-Consensus** | **70S-CBS** |
|---|---|---|---|
| Magnification | 105,000 | 105,000 | 105,000 |
| Voltage (kV) | 300 | 300 | 300 |
| Electron Exposure ($e^-/Å^2$) | 40 | 40 | 40 |
| Defocus Range (µm) | -0.5/-1.5 | -0.5/-1.5 | -0.5/-1.5 |
| Pixel Size (Å) | 0.8293 | 0.8293 | 0.8293 |
| Symmetry Imposed | C1 | C1 | C1 |
| Initial Particle Images | 2,360,632 | 2,360,632 | 2,360,632 |
| Final Particle Images | 183,467 | 129,829 | 72,157 |
| Map Resolution (Å) | 1.95 | 2.36 (Global) | 2.53 (Global) |
| FSC Threshold | 0.143 | 0.143 | 0.143 |

**Supplemental Table 7. Model Refinement Statistics**

| Model | **50S-CBS** | **70S-Consensus** | **70S-CBS** |
|---|---|---|---|
| Model component | | | |
| Model resolution (Å) | 2.07 | 2.52/2.90 (50S/30S) | 2.64/2.98 (50S/30S) |
| FSC threshold | 0.5 | 0.5 | 0.5 |
| Map sharpening *B* factor (Å$^2$) | 36.3 | 42.3/47.4 (50S/30S) | 41.8/43.8 (50S/30S) |
| Model composition | | | |
| Non-hydrogen atoms | 114588 | 171375 | 172200 |
| $Mg^{2+}$ ions | 173 | 252 | 251 |
| Waters | 7994 | 7812 | 6387 |
| Mean *B* Factors (Å$^2$) | | | |
| RNA | 49.06 | 66.88 | 66.12 |
| Protein | 56.83 | 70.87 | 72.98 |
| Waters | 48.77 | 60.66 | 61.99 |
| Other | 47.58 | 59.46 | 61.04 |
| R.m.s. deviations from ideal values | | | |
| Bond (Å) | 0.008 | 0.006 | 0.007 |
| Angle (°) | 0.891 | 0.898 | 0.907 |
| Molprobity score | 1.46 | 2.42 | 2.41 |
| Clash Score | 5.79 | 16.77 | 15.02 |
| Rotamer outliers (%) | 1.31 | 3.59 | 3.65 |
| Ramachandran plot | | | |
| Favored (%) | 97.74 | 96.02 | 95.67 |
| Allowed (%) | 2.23 | 3.72 | 4.08 |
| Outliers (%) | 0.04 | 0.26 | 0.24 |
| RNA validation | | | |
| Angles outliers (%) | 0.05 | 0.05 | 0.04 |
| Sugar pucker outliers (%) | 1.14 | 1.53 | 1.32 |
| Average suiteness | 0.62 | 0.568 | 0.575 |

**Supplemental Table 8. DNA primers used in this study**

| Primer Name | Sequence | Description |
|---|---|---|
| PTC_InFusion_F | GGGCGCTGGAGAACTGAGG | Forward primer for pLK35 linearization |
| PTC_InFusion_R | GGTACACTGCATCTTCACAGCGAGTTC | Reverse primer for pLK35 linearization |
| uL3_sgRNA_ins_F | cggtcagaaccagactccGTTTTAGAGCTAGAAATAGCAAG | Forward primer for uL3 N20 sequence insertion into the pEcgRNA plasmid with the InFusion kit |
| uL3_sgRNA_ins_R | gtctggttctgaccgatACTAGTATTATACCTAGGACTGAGC | Reverse primer for uL3 N20 sequence insertion into the pEcgRNA plasmid with the InFusion kit |
| uL3_HDR_seq_F | GGTTAATCAGGTCATTGAGCGATTGAGAGG | Forward primer for amplification of the *E. coli rplC* gene |
| uL3_HDR_seq_R | TTGACTGTGGTCCTGCGGACGAG | Reverse primer for amplification of the *E. coli rplC* gene |
| MS2_H98_quant_F | CTTGCCCCGAGATGAGTTCTCCC | RT-PCR primer for the *E. coli* ribosome purity assay with H98 MS2 tags |
| MS2_H98_quant_R | GTACCGGTTAGCTCAACGCATCGCT | Primer for amplification of cDNA in the *E. coli* ribosome purity assay with H98 MS2 tags |

**Supplemental Table 9. sgRNA sequence used in this study**

| Name | Sequence |
|---|---|
| uL3-ΔQ150 N20 | AUCGGUCAGAACCAGACUCC |
| uL3-ΔQ150 Full sgRNA | AUCGGUCAGAACCAGACUCCGUUUUAGAGCUAGAAAUAGCAAGUUAAAAUAA GGCUAGUCCGUUAUCAACUUGAAAAAGUGGCACCGAGUCGGUGC |

**Supplemental Table 10. dsDNA blocks used in this study**

| Name | Sequence |
|---|---|
| PTC_3M | AAGATGCAGTGTACCCGCGGCAAGACGGAAAGACCCCGTGAACCTTTACTATAGCTTGACACTGAACATTGAGCCTTGATGTGTAGGATAGGTGGGAGGCTTTGAAGTGTGGACGCCAGTCTGCATGGAGCCGACCTTGAAATACCACCCTTTAATGTTTGATGTTCTAACGTTGACCCGTAATCCGGGTTGCGGACAGTGTCTGGTGGGTAGTTTGACTGGGGCGGTCTCCTCCTAAAGAGTAACGGAGGAGCACGAAGGTTGGCTAATCCTGGTCGGACATCAGGAGGTTAGTGCAATGGCATAAGCCAGCTTGACTGCGAGCGTGACGGCGCGAGCAGGTGCGAAAGCAGGTCATAGTGATCCGGTGGTTCTGAATGGAAGGGCCATCGCTCAACGGATAAAAGGTACTCCGGGGATAACCGGCTGATACCGCCCAAGAGTTCATATCGACGGCGGTGTTTGGCACCCCGACCGTCGGCTCATCACATCCTGGGGCTGAAGTAGGTCCCAAGGGTATGGCTGTTCGCCATTTAAAGTGGTACGCGAGCTGGGTTTAGAACGTCGTGAGACAGTTCGGTCCCTATCTGCCGTGGGCGCTGGAGAACT |
| uL3-ΔQ150 HDR Template | GTTACTCAGGTTAAAGACCTGGCTAACGATGGCTACCGTGCTATTCAGGTGACCACCGGTGCTAAAAAAGCTAACCGTGTGACCAAGCCTGAAGCTGGCCACTTCGCTAAAGCTGGCGTAGAAGCTGGCCGTGGTCTGTGGGAATTCCGCCTGGCTGAAGGCGAAGAGTTCACTGTAGGTCAGAGCATTAGCGTTGAACTGTTTGCTGACGTTAAAAAAGTTGACGTAACTGGCACCTCTAAAGGTAAAGGTTTCGCAGGTACCGTTAAGCGCTGGAACTTCCGTACCCAGGACGCTACTCACGGTAACTCCTTGTCTCACCGCGTTCCGGGTTCTATCGGTCAGAACACTCCGGGCAAAGTGTTCAAAGGCAAGAAAATGGCAGGTCAGATGGGTAACGAACGTGTAACCGTTCAGAGCCTTGACGTAGTACGCGTTGACGCTGAGCGCAACCTGCTGCTGGTTAAAGGTGCTGTCCCGGGTGCAACCGGTAGCGACCTGATCGTTAAACCAGCTGTGAAGGCGTAAGGAGATAGCAATGGAATTAGTATTGAAAGACGCGCAGAGCGCGCTGACTGTTTCCGAAACTACCTTCGGTCGTGATTTCAACGAAGCGCTGGTTCACCAGGTTGTTGTTGCTTATGCAGCTGGTGCTCGTCAGGGTACTCGTGCTCAGAAGACTCGTGCTGAAGTAACTGGTT |
| HiBit DNA Template | GCGAATTAATACGACTCACTATAGGGTTAACTTTAACAAGGAGAAAAACATGGTGAGCGGCTGGCGCCTGTTTAAAAAAATTAGCTAACTAGCATAACCCCTCTCTAAACGGAGGGGTTT |
